# Role of *chrna5* in multi-substance preference and phenotypes comorbid with the development of substance dependence

**DOI:** 10.1101/2025.07.10.663858

**Authors:** Tanisha Goel, Joshua Raine, Caroline Kibat, Jeff Winxin Collado, Tirtha Das Banerjee, Ajay S. Mathuru

**Affiliations:** Department of Physiology, Yong Loo Lin School of Medicine, National University of Singapore, Singapore, Singapore; Yale-NUS College, National University of Singapore, Singapore, Singapore; Department of Biological Sciences, National University of Singapore, Singapore, Singapore; Institute for Digital Medicine (WisDM), Yong Loo Lin School of Medicine, National University of Singapore, Singapore; Institute of Molecular and Cell Biology, A*STAR, Singapore, Singapore; Healthy Longevity Translational Research Programme, Yong Loo Lin School of Medicine, National University of Singapore, Singapore, Singapore

**Keywords:** Zebrafish, Nicotine Dependence, Alcohol Use Disorders, SUD, Hormesis, *CHRNA5*

## Abstract

Addiction to nicotine and alcohol continues to be a leading cause of death and loss of productivity as measured in disability-adjusted life years. Polymorphisms in the nicotinic acetylcholine receptor subunit α5 (CHRNA5) have been identified as risk factors associated with nicotine dependence in human genetic studies and rodent models. Whether the *chrna5* function is independently relevant to phenotypes associated with disorders comorbid with substance use, and if genetic factors influence subsequent outcomes when exposure to psychoactive substances happens at an early age, are questions of interest. We generated a stable mutant line in zebrafish using the CRISPR-Cas9 technique. We found that the *chrna5* mutant fish exhibit an increased acute preference to both nicotine and alcohol in the Self-Administration Zebrafish Assay (SAZA). When subjected to multi-day exposures to either, *chrna5* mutants exhibited greater behavioural change, but reduced transcriptomic changes compared to WT siblings, suggesting an impaired homeostatic regulation following drug exposure. Further, *chrna5* mutants exhibited drug-independent changes in appetite and circadian rhythms, suggesting a genetic predisposition to disorders often comorbid with substance dependence. We expect these results to give new insights into the operation of genes whose normal function modulates vulnerability to multi-substance use and comorbid disorders.

## Introduction

Substance Use Disorders (SUDs), due to the abuse of nicotine, alcohol, and opioids, are a major global health burden today, with an estimated 162.5 million disability-adjusted life years lost to them globally in 2016 [1–3]. The direct death toll from tobacco alone is predicted to exceed 8 million per year by 2030 [4,5], with cancers, cardiovascular diseases, and chronic obstructive pulmonary disorders as the major disease outcomes [6]. The development of dependence leading to these outcomes is influenced by a complex mix of socio-economic, cultural, neurobiological, and genetic factors that is challenging to untangle [7,8].

A particularly vulnerable period of substance exposure is early age, when the still-developing brain undergoes significant neurodevelopmental and neuroanatomical changes. Exposure during this critical window can lead to profound alterations in brain structure and gene expression that can be long-lasting [9,10]. Genetic variation contributes to differences in individual susceptibility, both for the consequences of exposure at an early age and the risk of developing SUDs subsequently [9]. Research aimed towards isolating the genetic components has revealed strong associations between polymorphisms in nicotinic acetylcholine receptor subunit genes (nAChRs) and nicotine use [8]. In particular, the gene cluster *CHRNA5-CHRNA3-CHRNB4* encoding for the α5, α3, and β4 subunits has been frequently identified in genome-wide association studies [11–16].

In rodents, where *Chrna5* function has been examined extensively [17], expression across many regions of the brain, including medial habenula-IPN pathway [18] and ventral tegmental area (VTA [19]) is documented. These regions are proposed to impact withdrawal behaviours and nicotine dependence [20]. Manipulation of *Chrna5* levels within them has been shown to alter nicotine preference [18,21]. In addition to nicotine, polymorphisms in the *CHRNA5-CHRNA3-CHRNB4* locus have also been correlated with vulnerability to alcohol dependence [22,23], despite alcohol not being a direct ligand [23–26]. As co-abuse of nicotine and alcohol is common [27], elucidation of the interdependent relationships represents an important step towards conceiving holistic interventions. Multi-day exposure to nicotine and alcohol impacts both the transcriptional profile and behaviour, and it is equally important to understand the neuroadaptive changes in the brain that influence behaviour [28–30]. Furthermore, the harmful outcomes of SUDs are not limited to addiction, as neurophysiological disorders of anxiety, sleep, and appetite control often occur comorbidly [31]. Therefore, understanding the mechanisms underlying the initial stages of dependence, at an age when brains are still developing, is crucial for the design of interventional strategies informed by genetic predisposition.

Untangling the contribution of genetic predisposition to induction of these comorbid disorders is challenging to discover in humans, however, due to the bidirectional nature of the relationships between drug use and neuroplasticity, and ethical considerations, such as assigning an adolescent to drug exposure in a randomized control trial. In this context, animal models can be effective intermediates to bridge the gap in our knowledge. In addition to the mammalian models, the zebrafish represents a viable alternative to study neurogenetics, thanks to the established conservation of function and anatomy [32–34], cost effectiveness, efficient genetic manipulation, and live, whole brain neural activity imaging [35]. At the same time, a battery of behavioural assays to rapidly examine anxiety-like behaviours [36], circadian rhythms [37], and appetite [38] are now available. New assays to examine not just the effect of psychoactive substances [39], but also the natural responses [40–43] add to their value for neurogenetic studies.

Here, we generated a zebrafish *chrna5* mutant line using CRISPR-Cas9 technique to study transcriptomic and behavioural change after exposure to substances of abuse at an early age. In drug preference assays, homozygous *chrna5* mutant juvenile fish phenocopied adult rodent responses, exhibiting increased acute, naive self-administration of both nicotine and alcohol. This recapitulation validates the use of this model system to study the neurogenetics of development of SUDs. Multiday, pre-exposure experiments revealed the modulation of preference, with greater influence on *chrna5* mutant behaviour, while homeostatic transcriptomic changes reduced behavioural changes in WT animals. *chrna5* mutants were also impaired in circadian rhythm and appetite regulation, with no effect on anxiety-like behaviours. This study thus adds to our knowledge of α5 nicotinic acetylcholine receptors in the development of SUDs and the potential consequences of manipulating its function as an intervention.

## Results

The *chrna5* mutant zebrafish generated for this study by CRISPR-Cas9 had a five-base pair deletion in exon 8, resulting in a premature termination codon at amino acid position 334 (Figure 1(**a**)). This stop codon was predicted to truncate the protein at the intracellular loop between the third and fourth transmembrane regions (Figure 1(**b**)). In addition to the functional inhibition stemming from the introduction of a premature termination codon, we also examined the stability of the expression profile by quantitative RT-PCR of the adult whole brain. This examination indicated a small, but non-significant, reduction in the expression of *chrna5* transcripts in the mutants (Figure 1(**c**), WT vs. *chrna5*, Cliff’s delta = -0.438, 95CI [0.156, -0.875], *p* = 0.122). The reduction of *chrna5* mRNA expression resulted in a corresponding downstream reduction in the relative abundance of chrna5 protein levels in adult brains (Figure S5). The expression of *chrna3* and *chrnb4* was unchanged (Figure S1). *Chrna5* expression in mice has been reported in the habenula-interpeduncular nucleus (IPN) circuit in addition to the ventral tegmental area (VTA), relevant to nicotine aversive responses [17,45]. In zebrafish, *chrna5* mRNA expression has been reported in the ventral IPN as well [46], while chrna5 protein in a recently established transgenic line has been reported to be broader, in the pineal gland, stratum periventricular of the optic tectum, corpus cerebellum, and hindbrain motor neurons [47]. We examined the expression of zebrafish *chrna5* in 14 dpf larvae using RAM-FISH method developed in-house [48]. This analysis revealed that neurons in the telencephalon, torus longitudinalis, cerebellum, and hindbrain exhibited the highest expression, but low levels of *chrna5* expression were visible in the majority of brain regions, including the dorsal habenula (Figure 1(**d**) and S2-4).

**Figure 1.**
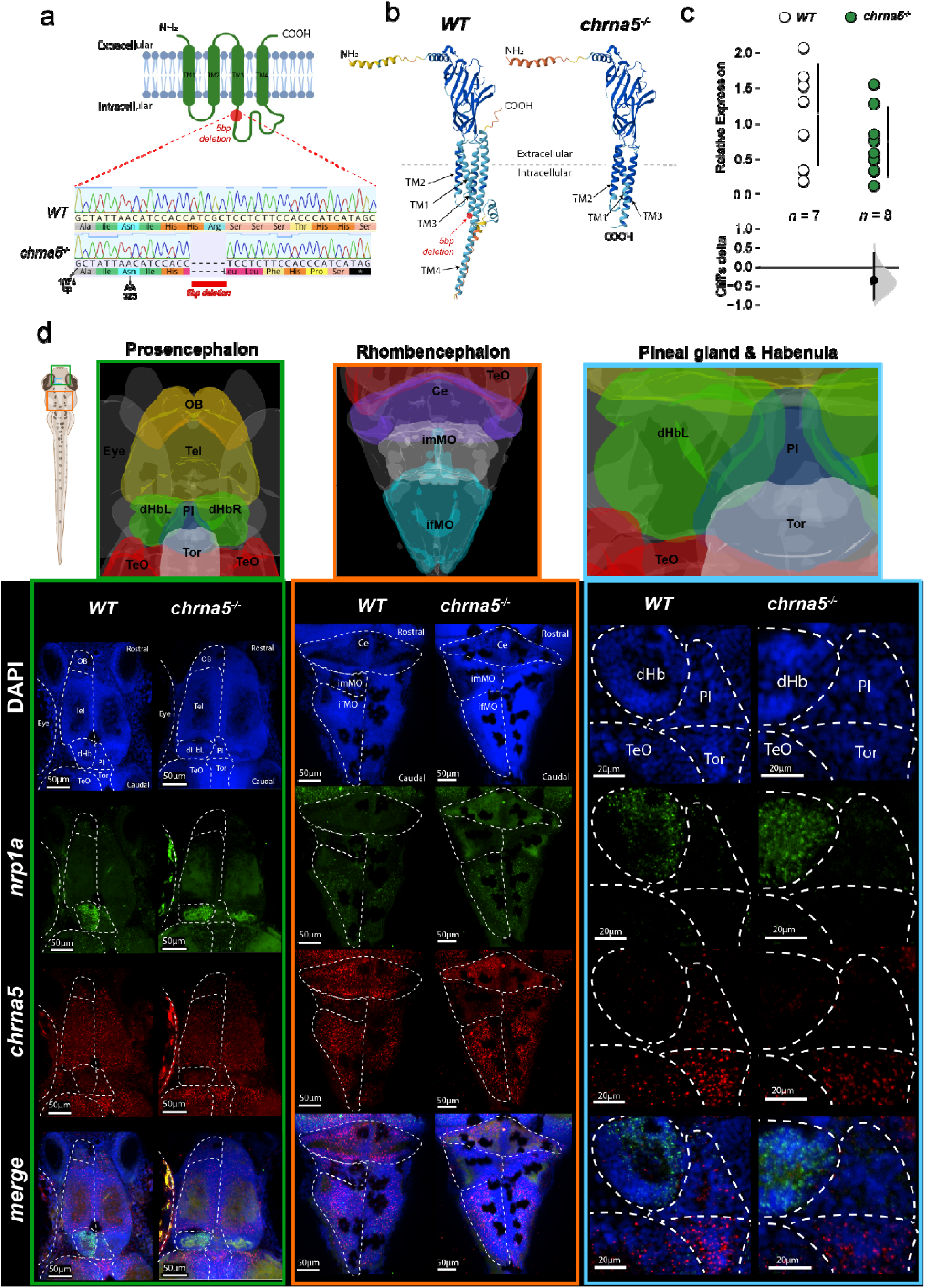
Characterisation of *chrna5* mutant zebrafish. (**a**) Schematic representation of chrna5 transmembrane helices. The red line indicates the stop codon present in the mutants between the third and fourth transmembrane regions. Mutants had a 5 bp deletion in exon 8 at position 10, resulting in a premature termination codon at amino acid 334 (UniProt entry Q567Y7_DANRE). (**b**) Predicted tertiary protein structure of chrna5 in WT and *chrna5* mutants, generated by Alphafold [44]. (**c**) Fold change of *chrna5* mRNA in WT (n = 7) vs. *chrna5* mutants (n = 8) whole brain tissue by qPCR. See table S3 for the precise effect size and p-values. (**d**) Gene expression patterns in 14 dpf larval zebrafish brains visualised by HCR™ RNA-FISH (HCR). Schematic images with regional markers (v2.0, MECE, 2024) were generated in mapZeBrain Atlas (mapzebrain.org, Jan 2025) (Kunst et al., 2019). Hb = habenula, vHb = ventral habenula, Ipn = interpeduncular nucleus, Omn = oculomotor nucleus, Trm = trochlear motor nucleus, Ni = nucleus isthmi, Vmn = vagus motor nucleus, srFmn = supra-rostral Fmn, rFmn = rostral Fmn, cFmn = caudal Fmn.

Given the strong links reported between loss-of-function *Chrna5* mutations, or knockouts, and altered intake of nicotine and alcohol in rodent studies [18,21,24], we first characterised if zebrafish *chrna5* mutants also showed similar changes in spite of the evolutionary distance to mammals. To do so, we used the self-administration for zebrafish assay (SAZA) previously reported [40,41] to examine the acute preference to self-administer nicotine and alcohol. Briefly, this assay allowed juvenile zebrafish an uninhibited choice to self-administer a stimulus, conditional on the fish’s behaviour of swimming into a dispensing region of the tank (Figure 2(**a**)). In the assay, a three minute pre and a post-stimulus periods were interspersed with an eighteen minute self administration period (Figure 2(**a**)). The parameters, including the volume of stimulus or control dispensed, were collected to calculate the relatve preference and absolute behaviour of the animals. The dispensing concentratons of 10/500μM nicotne and 5/10/20% alcohol were diluted to estmated mean concentratons of 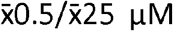 nicotne and 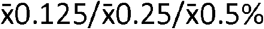 alcohol in the stimulus zone, as outlined in Table S2. These terms are used for the remainder of the work.

**Figure 2.**
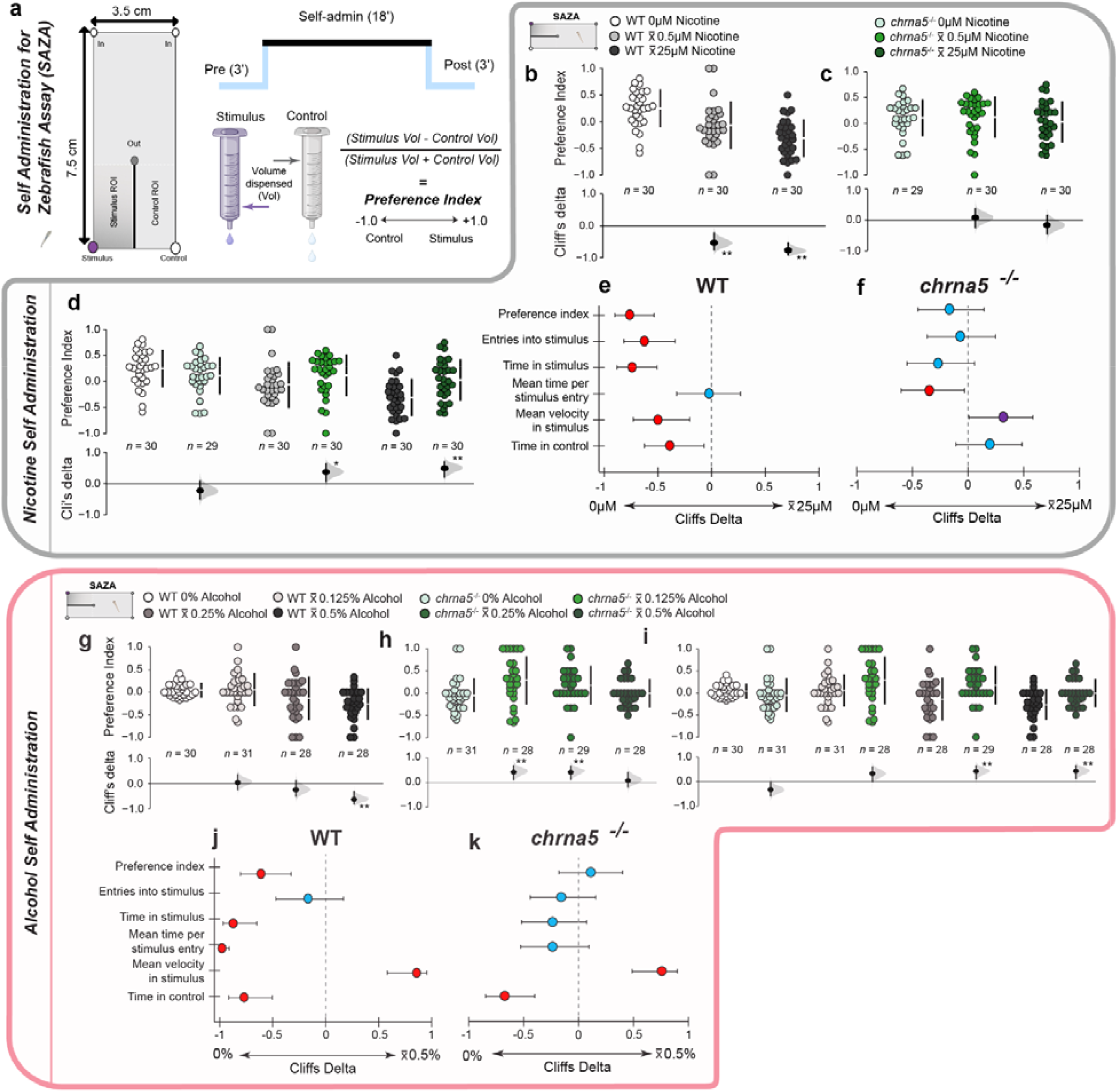
Zebrafish with *chrna5* mutation exhibited reduced aversion to acute nicotine & alcohol exposure by SAZA. (**a**) Schematic of the self administration for zebrafish assay (SAZA) chamber and design. (**b-d, g-i**) Gardener-Altman and Cumming estimation plot dose response curves displaying preference index by relative volume dispensed per fish of (**b-d**) 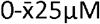 nicotine, or (**g-i**) alcohol, within (**b, g**) WT (greys), (**c, h**) *chrna5* mutant (greens), or (**d**,**i**) between genotypes. Positive values indicate preference for the stimulus. (**e-f, j-k**) Cliff’s delta (± 95% CI) forest plots of all calculated metrics from (**e, f**) nicotine and (**j, k)** alcohol SAZA, comparing the response of (**e, j**) WT or (**f, k**) c*hrna5* mutants to their baseline behaviour within genotype. Positive values indicate a greater response in the mutant. Asterisks (**b-d, g-i**), or colour (**e-f, j-k**) indicate a significant difference: /blue = p > 0.05 (no significant difference), */purple = p < 0.05 and Cliff’s delta > ± 0.2 & < ± 0.4 (provisional difference), **/red = p < 0.01 and Cliff’s delta > ± 0.4 (meaningful difference). See tables S4-5 for the precise effect sizes and p-values, corrected for multiple comparisons.

Wild-type fish (WT) exhibited a strong aversion to nicotine, with increasing magnitude of avoidance correlated with dispensing concentration (Figure 2(**b**)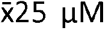, Cliff’s Delta = -0.762, 95%CI [-0.900, - 0.533], *p* < 0.01). However, this aversive response was blunted across the concentrations in *chrna5* mutants, with the preference index of neither the 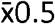 nor 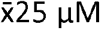 nicotine differing from the behaviour of the fish when no nicotine (0 μM) was administered (Figure 2(**c**). This resulted in a reduced aversion to nicotne in the mutant that was apparent at both concentratons, with the largest effect observed at 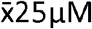 nicotine (Figure 2(**d**), 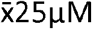, Cliff’s Delta = -0.166, 95%CI [-0.488, 0.143], *p* = 0.2688). Overall, these results indicated that the mutation affected the phenotype at higher concentrations of nicotine self-administration, reducing the impact on aversion.

Alcohol also evoked an aversive response in WT fish, albeit to a lesser degree than nicotne, exhibitng a significant change only under the 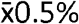 condition (Figure 2(**g**)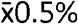, Cliff’s Delta = -0.611, 95%CI [- 0.810, -0.327], *p* < 0.01). When mutant fish were exposed to alcohol, however, both the 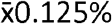 and 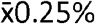 conditions evoked an attraction (Figure 2(**h**) 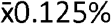, Cliff’s Delta = 0.447, 95%CI [0.121, 0.691], *p* = 0.0068, 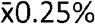, 95%CI [0.131, 0.665], *p* = 0.0018). In addition, as with nicotine, the response to the aversive 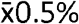 condition was also blunted in the mutants (Figure 2(**h**) 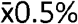, Cliff’s Delta = -0.109, 95%CI [-0.181,0.399], *p* = 0.4654). Again, the inter-genotype comparisons highlighted larger effect sizes and a more defined phenotype at the higher concentrations (Figure 2 (*i* 0.5%, Cliff’s Delta = 0.430, 95%CI [0.135,0.667], *p* = 0.0046). A breakdown of time spent in the stimulus into three-minute windows during the administration period at the highest concentrations of nicotine and alcohol revealed that the behaviour of the mutant fish was driven by a delay in the onset of an aversive response in comparison to the WT (Figure S6). Consistent with the behaviour observed in response to alcohol, the *chrna5* mutants were not tolerant to high concentrations of nicotine either and exhibited an aversion in the later stages of SAZA (Figure S6). The difference between the WT and *chrna5* mutant fish behaviour was negligible in the absence of nicotine or alcohol, indicating that the *chrna5* mutation did not interfere with normal behaviour in the SAZA apparatus (Figure 2(**d, i**). In the presence of either nicotine or alcohol, the evaluation of other behavioural metrics supported the observation of blunted aversion/increased tolerance to these substances (Figure 2 (**e-f, j-k** 25µM nicotine administration conditions, WT fish greatly reduced their time spent in the stimulus zone and number of entries compared to their behaviour in the absence of nicotine, while *chrna5* mutants exhibited no change in these measures (Figure 2 (**e**-**f** 0.5% alcohol SAZA, WTs reduced their time spent in the stimulus zone and mean time per entry versus baseline, while mutants showed no change (Figure 2 (**j-k**)). In all cases, changes in mean velocity of the fish were observed, a strong indicator for the stimuli delivered in the assay being experienced by the subjects, and altering their preference behaviours (Figure 2 (**j-k**)). Therefore, under these acute exposure parameters, *chrna5* mutants exhibited a phenotype of increased tolerance to substances of abuse, much like those reported in rodents.

We next evaluated the impact of the genetic predisposition on the process of development of substance dependence. To do so, we examined the effect of multi-day exposures on self-administration preference, as substance abuse happens after recurring exposure to these stimuli that alter both gene expression and behaviour [29,49]. Further, nicotine addiction and alcohol abuse are often comorbid in humans [50,51]. Towards this end, we designed a substance pre-exposure scheme before examining gene expression of the forebrain and midbrain (excluding the olfactory bulb, telencephalon, and hindbrain regions) using bulk RNA-seq and the behavioural response of subject fish in SAZA. Both WT and *chrna5* mutant fish were subjected to a one-week pre-treatment to nicotine or alcohol prior to SAZA, and the SAZA to both substances was conducted following this scheme (Figure 3).

**Figure 3.**
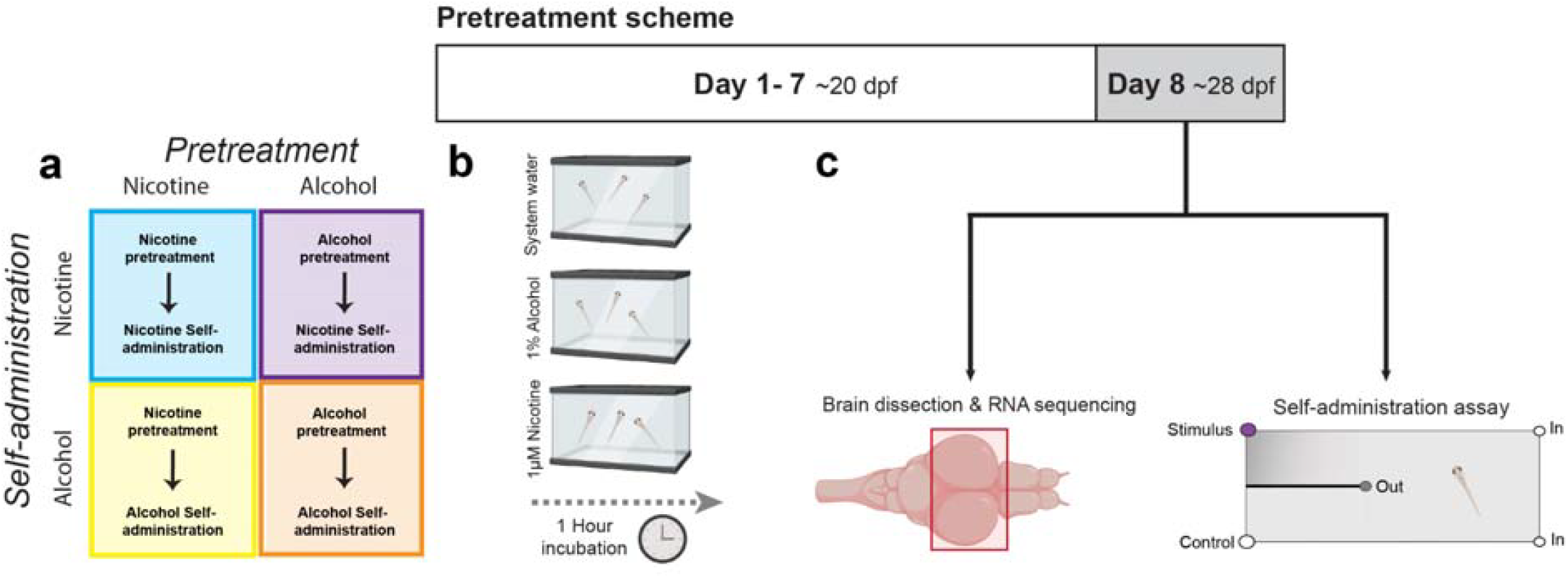
Combined multi substance behavioural and transcriptomic analysis schematic detailing **(a)** pre-treatment and self-administration combinations, **(b)** the pre-treatment schemes for alcohol and nicotine repeated daily over seven days, and **(c)**, the analyses performed on the pre-treated fish.

Pre-treatment of the WT fish with nicotine appeared to have little impact on their behavioural profile for self-administering nicotine (Figure 4(**a**), 2(**e**)), or alcohol (Figure 4(**d**), 2(**j**)). PI, time in stimulus, and number of entries to the stimulus zone all exhibited drastic reductions compared to baseline behaviours (Figure 4(**a**) PI, Cliff’s Delta = -0.792, 95%CI [-0.917,-0.572], *p* < 0.01), as was the case with the non-pre-treated fish (Figure 2(**e**)). Unlike their untreated counterparts, *chrna5* mutants however, exhibited a small but significant negative shif in preference index and tme spent in stimulus in the 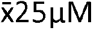 nicotine SAZA following nicotine pre-treatment (Figure 4(**b**) PI, Cliff’s Delta = -0.414, 95%CI [-0.667, -0.0989], *p* = 0.0058), suggesting a reduced tolerance compared to the untreated phenotype (Figure 2(**f**)). This narrowed the phenotypic gap between the two genotypes slightly (Figure 4(**c**) pre-treated mutant vs WT, Cliff’s Delta = 0.338, 95%CI [0.0258, 0.578], *p =* 0.0244), though whether this was due to a true lack of change in phenotype in the WT fish, or if the lower bounds to quantify avoidance behaviours in the SAZA had already been reached, require additional lines of experimentation.

**Figure 4.**
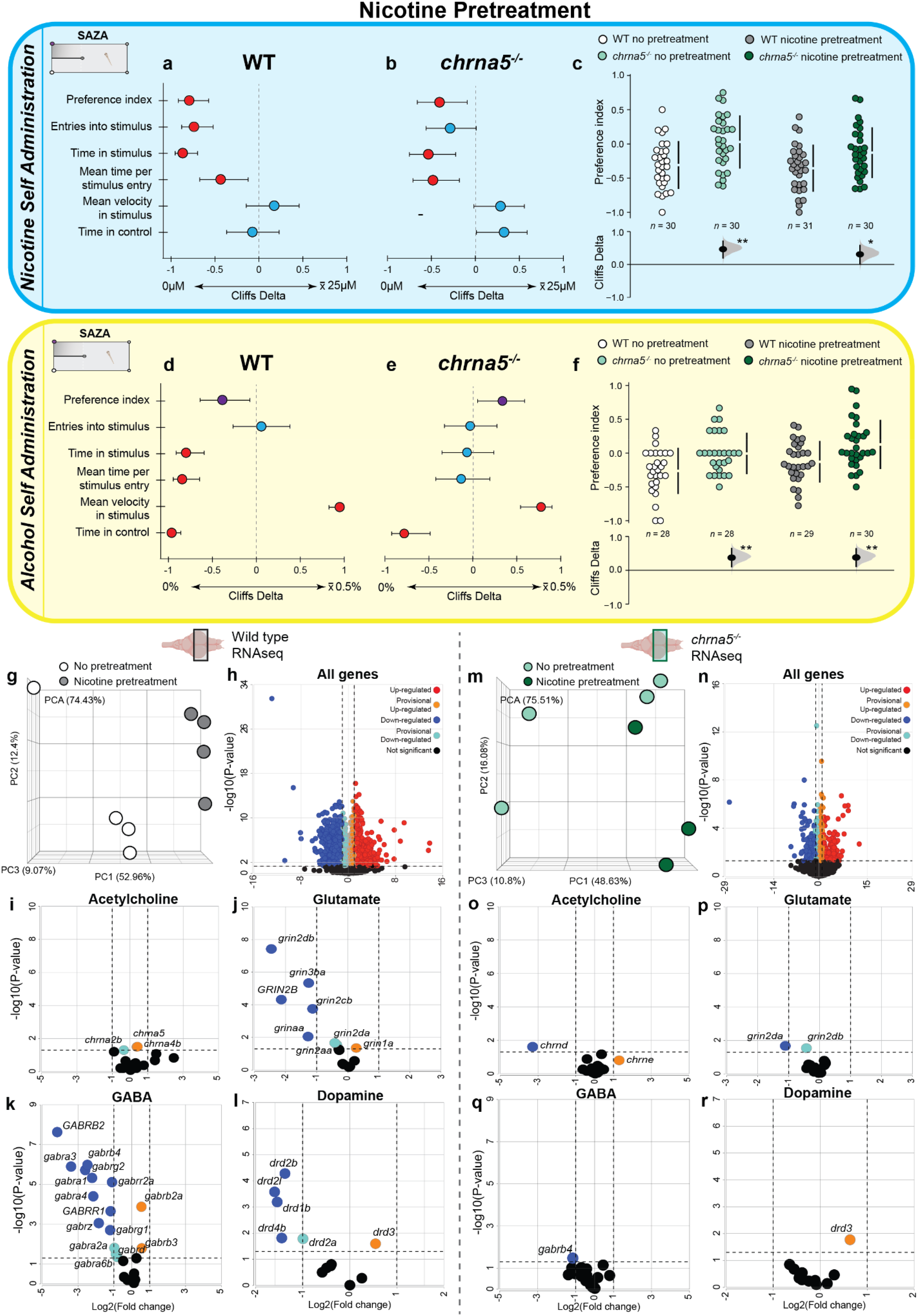
Nicotine pre-treatment altered *chrna5* mutant acute behavioural responses to nicotine and alcohol, and effected transcriptomic changes in WT fish. (**a-f**) Measures of fish behaviour in SAZA following a seven day nicotine pre-treatment scheme when self-administering (**a-c**) 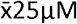 nicotine or (**d-f**) 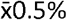 alcohol. Comparisons of multiple measures in (**a, d**) WT or (**b, e**) *chrna5* mutants to baseline, or (**c**,**f**) to each other by preference index, are displayed as forest plots, or Gardener-Altman and Cumming estimation plots, respectively. Asterisks (**c, f**), or colour (**a-b, d-e**) indicate a significant difference: blue = p > 0.05 (no significant difference), */purple = p < 0.05 and Cliff’s delta > ± 0.2 & < ± 0.4 (provisional difference), **/red = p < 0.01 and Cliff’s delta > ± 0.4 (meaningful difference). See table S6 for the precise effect sizes and p-values, corrected for multiple comparisons. (**g-r**) RNA sequencing data of (**g-l**) WT and (**m-r**) *chrna5* mutant fish brain tissues, with the olfactory bulb, telencephalon and hindbrain removed, comparing untreated samples to those collected following a seven day nicotine pre-treatment scheme. (**g**,**m**) PCA separation of samples. (**h-l, n-r**) Volcano plots of (**h**,**n**) all genes, (**i**,**o**) nicotinic acetylcholine receptors (chrn), (**j**,**p**) glutamate receptors (grin), (**k**,**q**) GABA receptors (gabr), (**l**,**r**) dopamine receptors (drd). Significance was categorised as adj *p* > 0.05 = non significant (black), adj *p* < 0.05, log_2_ fold change 0 - ±1 = provisional up (+, orange) or down (−, cyan) regulation, adj *p* < 0.05, log_2_ fold change ±1 - >±2 = up (+, red) or down (−, blue) regulation.

Unlike in nicotine self-administration, the behavioural profile of WT fish to self-administering alcohol in the SAZA assay indicated reduced aversion following nicotine pre-treatment, compared to their untreated counterparts (Figure 4(**d**) PI, Cliff’s Delta = -0.383, 95%CI [-0.637, -0.0713], *p* = 0.011). The *chrna5* mutant fish also exhibited a higher preference for 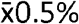 alcohol following nicotine pre-treatment (Figure 4(**e**) PI, Cliff’s Delta = 0.335, 95%CI [0.0516, 0.587], *p* = 0.0252). This is in contrast to the response of untreated mutants, which displayed a neutral preference (Figure 2(**k**)). This suggests that nicotine pre-treatment amplified the alcohol tolerance phenotype previously observed for the mutants and WT alike. Next, we performed bulk RNA-sequencing on the midbrains of fish that had undergone the nicotine pre-treatment to get a deeper understanding of potential causes for these behavioural shifts from the transcriptomes. While untreated and pre-treated samples were separated in the PCA for both genotypes (Figure 4 (**g, m**)), nicotine pre-treatment of c*hrna5* mutants appeared to effect fewer transcriptomic changes than in WT in the dissected brain regions (Figure 4 (**h, n**)). Nicotinic acetylcholine receptor subunit expression was unchanged after the pre-treatment in either genotype (Figure 4 (**i, o**). Interestingly, several genes in the glutamate, GABA, and dopamine receptor families were significantly downregulated only in the WT fish (Figure 4 (**j-l, p-r**)) that were unaltered in the mutants. Untreated WT and *chrna5* mutant fish showed few transcriptomic differences (Figures S8-9).

We next examined the self-administration behaviour of fish pre-treated with alcohol. Similar to the nicotine pre-treatment results, there was no observable impact on the self-administration behaviours of WT fish. WT fish continued to avoid 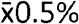 alcohol (Figure 5(**a**) PI, Cliff’s Delta = -0.617, 95%CI [-0.799, - 0.347], *p* < 0), similar to the strong aversion seen without pre-treatment (Figure 2(**j**)). Mutants, in this case, were unchanged in their behavioural measures to alcohol self-administration and resembled untreated mutant fish (Figure 5(**b**), 2(**k**)). The only notable difference between the alcohol pre-treated and untreated fish being time per entry (Figure 5(**b**) mean time per entry, Cliff’s Delta = 0.5, 95%CI [0.181, 0.739], *p* = 0.004). Inter genotype comparison, however, revealed that the difference observed between untreated WT and mutants had been abolished following the alcohol pre-treatment (Figure 5(**c**) WT vs mutant pre-treated, Cliff’s Delta = 0.225, 95%CI [-0.0927, 0.504], *p* = 0.1302). This appeared to be due to an increase in variance amongst the assayed mutant population, which exhibited a more polar distribution.

**Figure 5.**
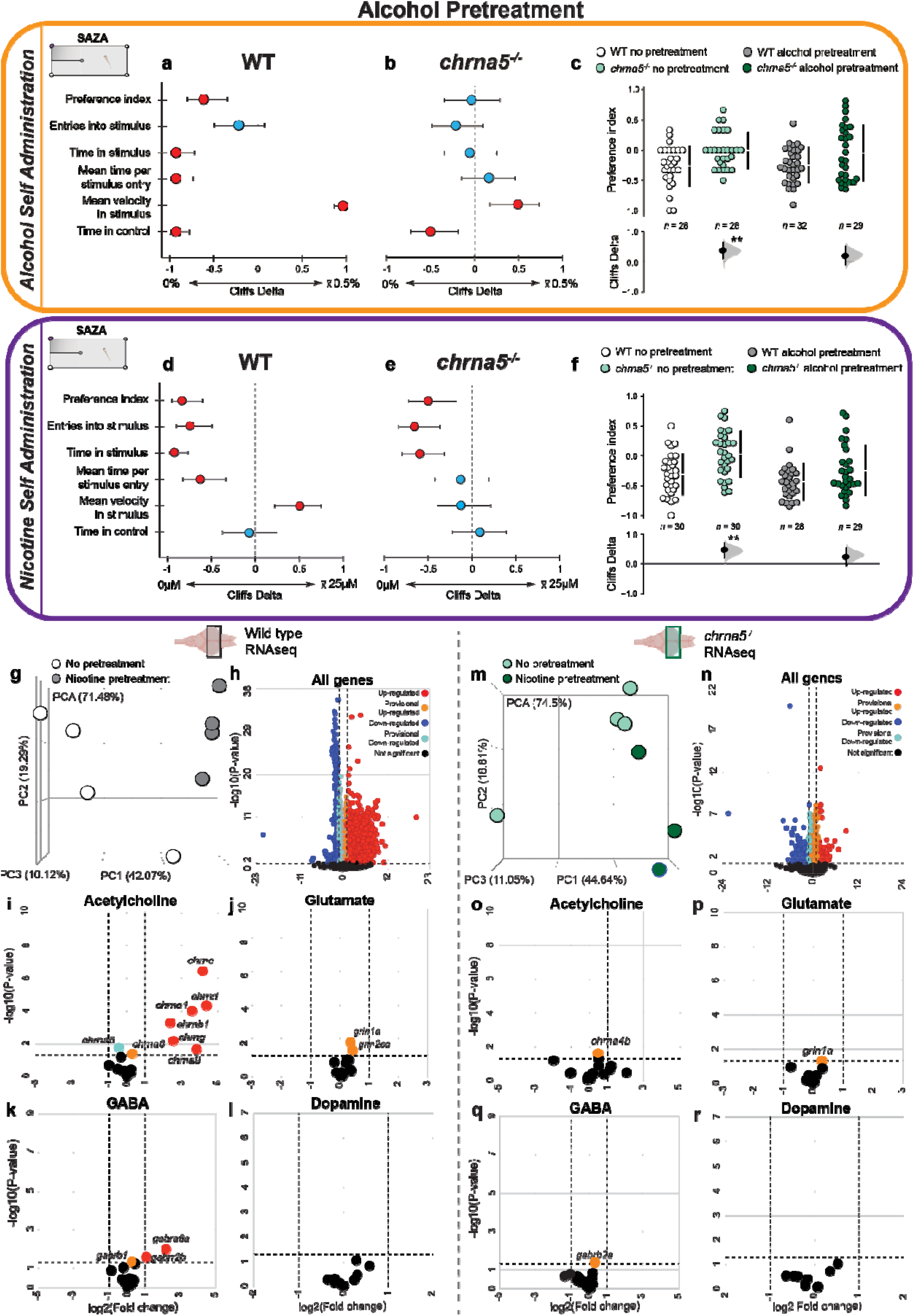
Alcohol pre-treatment abolished *chrna5* mutant nicotine tolerance phenotype, and stimulated upregulation of WT nicotinic acetylcholine receptor genes (**a-f**) Measures of fish behaviour in SAZA following a seven day alcohol pre-treatment scheme when self-administering (**a-c**) 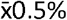 alcohol or (**d-f**) 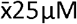 nicotine. Comparisons of multiple measures in (**a, d**) WT or (**b, e**) *chrna5* mutants to baseline, or (**c, f**) to each other by preference index, are displayed as forest plots, or Gardener-Altman and Cumming estimation plots, respectively. Asterisks (**c, f**), or colour (**a-b, d-e**) indicate a significant difference: blue = p > 0.05 (no significant difference), */purple = p < 0.05 and Cliff’s delta > ± 0.2 & < ± 0.4 (provisional difference), **/red = p < 0.01 and Cliff’s delta > ± 0.4 (meaningful difference). See table S7 for the precise effect sizes and p-values, corrected for multiple comparisons. (**g-r**) RNA sequencing data of (**g-l**) WT and (**m-r**) *chrna5* mutant fish midbrain tissues comparing untreated samples to those collected following a seven day alcohol pre-treatment scheme. (**g, m**) PCA separation of samples. (**h-l, n-r**) Volcano plots of (**h, n**) all genes, (**i, o**) nicotinic acetylcholine receptors (chrn), (**j**,**p**) glutamate receptors (grin), (**k, q**) GABA receptors (gabr), (**l, r**) dopamine receptors (drd). Significance was categorised as adj *p* > 0.05 = non significant (black), adj *p* < 0.05, log_2_ fold change 0 - ±1 = provisional up (+, orange) or down (−, cyan) regulation, adj *p* < 0.05, log_2_ fold change ±1 - >±2 = up (+, red) or down (−, blue) regulation.

Finally, we examined the effects of cross treatment on self administration here as well. Alcohol pre-treated fish were given the choice to self administered 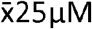 nicotine. WT fish in this condition showed aversive behaviour broadly similar to their untreated counterparts (Figure 5(**d**)). The *chrna5* mutants were examined next. Here too, as was the case with the nicotine pre-treated fish, alcohol pre-treatment appeared to reduce the tolerance of mutants to nicotine. Nicotine self-administration across multiple measures, including PI changed towards avoidance (Figure 5(**e**) PI, Cliff’s Delta = -0.502, 95%CI [-0.731,-0.182], *p* < 0). This increase in aversive behaviours resulted in a narrowing of phenotypic differences originally seen between the genotypes. The PI difference was no longer significant (Figure 5(**f**) WT vs mutant pre-treated, Cliff’s Delta = 0.260, 95%CI [-0.0603, 0.527], *p* < 0). RNA sequencing of the midbrains of the alcohol pre-treated fish revealed a phenomenon parallel to that observed after nicotine pre-treatment. The pre-treated and untreated samples of each genotype were separated by PCA (Figure 5 (**g, m**)), but again the scale of transcriptomic changes in the *chrna5* mutants amongst all genes was sharply lower than the transcriptome changes in the WT (Figure 5 (**h, n**)). Curiously, the pattern of expression changes amongst the neurotransmitter systems was almost the complete inverse of that observed after nicotine pre-treatment. Alcohol pre-treatment stimulated strong upregulation amongst many nicotinic acetylcholine receptor subunits (Figure 5(**i**)), but little change was observed in the glutamate, GABA, and dopamine receptors that were downregulated after nicotine pre-treatment. Once again, *chrna5* mutants exhibited little alteration to the transcriptomic profiles of these receptor genes (Figure 5 (**o-r**)). Thus, although pre-treatment with nicotine or alcohol had no notable population level change in the self-administration behaviours of WT fish, it resulted in large scale gene expression changes within the dissected brain regions. On the other hand, the behaviour of the mutant *chrna5* fish changed with limited changes in transcriptional profiles of the midbrain.

Nicotine dependence is frequently comorbid with a variety of disorders, including increased anxiety, alteration in circadian rhythms, and appetite dysregulation in humans [52–54]. While specific variants in the CHRNA5-CHRNA3-CHRNB4 gene cluster, particularly the CHRNA5 variant rs16969968, are strongly associated with nicotine dependence [55,56], a direct link between CHRNA5 and comorbid disorders more tentative [57]. However, a genetic predisposition for comorbid disorders is proposed [20]. Given that our *chrna5* mutant zebrafish exhibited a tolerance phenotype towards nicotine and alcohol that could act as a pathway to dependence, we evaluated if the mutation impacts any other behaviours in the fish. Nicotine and alcohol are both described as drugs that can have an anxiety reducing effect at specific doses in both humans and zebrafish [58–61]. As such, the drive to ameliorate anxiety is cited as a potential cause for the use of these psychoactive drugs. Whether genetic predisposition to developing drug dependence also causes individuals to have an anxiety phenotype due to a gene mutation or variant is debated [20]. To determine if *chrna5* mutant zebrafish also exhibit higher anxiety-like behaviours, we used a light/dark assay. 12-14 dpf zebrafish naturally exhibit scotophobia, or dark avoidance behaviours [62]. The behaviour of fish that were allowed to swim uninhibited between two zones, one illuminated and one darkened, was quantified (Figure 6(**a**)). Individuals spending more time in the dark, entering the dark sooner or more frequently, are all representative of a state of reduced anxiety-like behaviour in larval zebrafish. Both WT and mutant fish spent more time in the illuminated areas and entered the darkened area only after a few minutes. In all of these measures (Figure 6 (**b-d**), S7) no significant difference was observable between the WT and *chrna5* mutants. Thus, *chrna5* had little to no direct involvement in governing anxiety-like phenotypes in fish.

**Figure 6.**
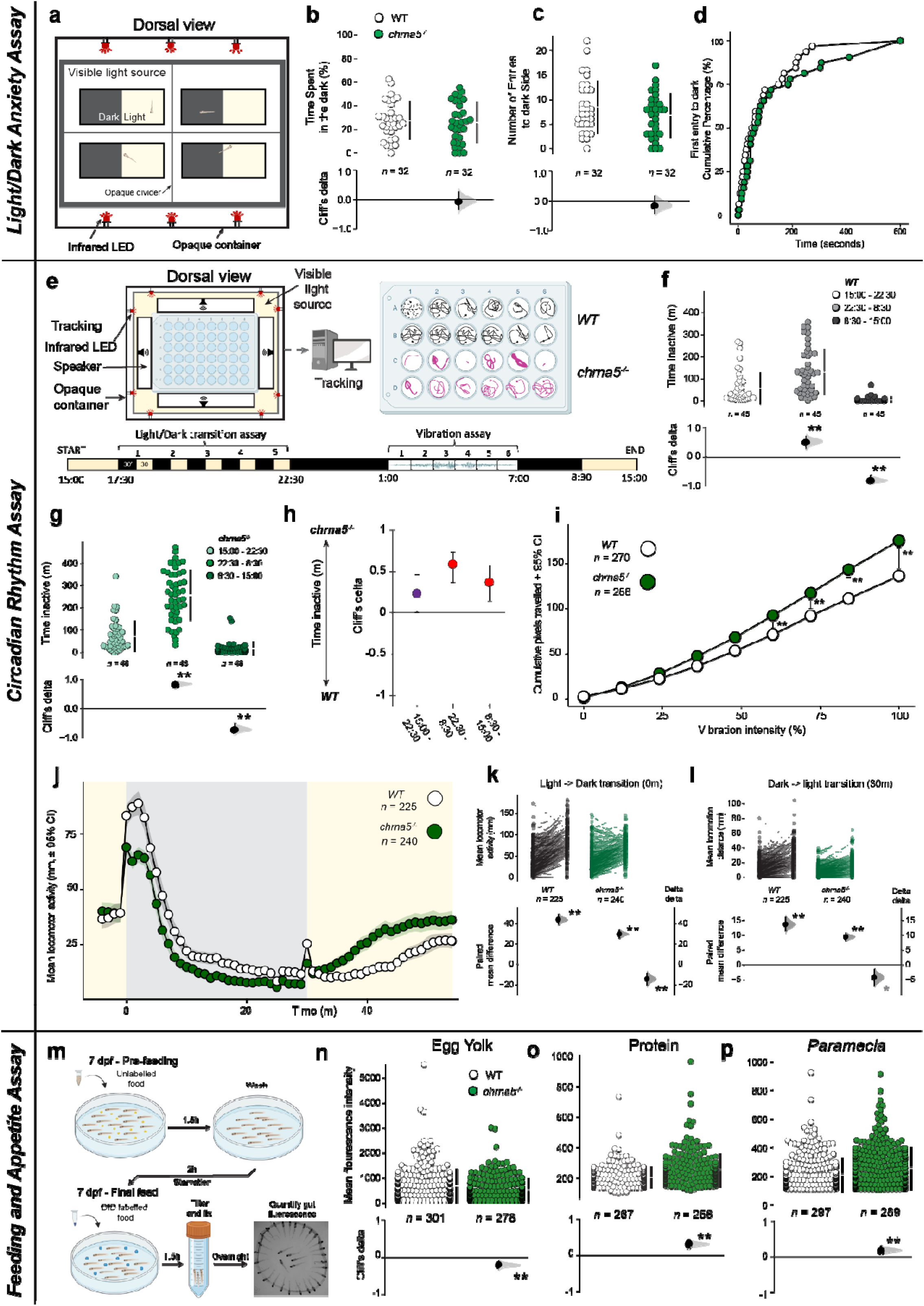
Appetite and circadian rhythm disorder associated phenotypes were altered in *chrna5* mutants. **(a-d).** Anxiety-like behaviour of 12-14 dpf larvae in light/dark assay. **(a)** Schematic of the light/dark assay equipment. Chambers were illuminated from below, divided into equal light and dark halves, and movement between the areas was tracked. **(b-d)** Behavioural measures of **(b)** time spent in dark, **(c)** number of entries to dark, and **(d)** cumulative first entry time of the population to the dark over the assay duration between WT and *chrna5* mutant fish. **(e-l)** Circadian rhythm, vibration sensitivity, and light-dark transition behaviours of 7-10 dpf fish. **(e)** Schematic of the circadian assay equipment and tracking (upper), and the assay timeline (lower). Yellow indicates visible lights are on, while black is off. **(f-h)** Time inactive over assay duration, divided according to light on/off times in **(f)** WT, **(g)** c*hrna5* mutants, and **(h)**, between genotypes. **(i)** Mean cumulative distance moved by fish per vibration stimulus intensity between genotypes. **(j-l)** Light-dark transition locomotor response. **(j)** Mean locomotor activity across all five cycles between genotypes. Yellow/grey areas indicate visible light on and off, respectively. **(k-l)** Difference in individual locomotor activity within and between genotypes (delta-delta) from before and after **(k)** light to dark transition at 0 minutes, and **(l)** dark to light transition at 30 minutes. **(m-p)** Feeding behaviour of 7-dpf fish given assorted food types. **(m)** Schematic detailing feeding protocol, pre-feeding, and starvation periods prior to the labelled feed. **(n-p)** Mean fluorescent intensity gut intensity of WT and *chrna5* mutant fish when fed **(n)** egg yolk, **(o)** protein, and **(p)** *paramecia*. **\* =** *p* < 0.05 and Cliff’s Delta from ±0.2 to ±0.4 (where applicable), ** **=** *p* < 0.01 and Cliff’s Delta from > ±0.4 (where applicable). See tables S8-9 for the precise effect sizes and p-values, corrected for multiple comparisons.

Nicotine, along with other substances of abuse, has also been implicated in the disruption of sleep and circadian rhythms both directly in animal models, and by comorbidity in human genetic studies [63–66]. However, despite the nicotinic acetylcholine receptor family being involved in the maintenance of circadian rhythms and the sleep-wake cycle [67], whether *chrna5* plays a role is untested. To study the circadian rhythm, we tracked the activity of 7-10dpf larval zebrafish over 24 hours, starting in the afternoon (Figure 6(**e**)). Over a period of 24 hours, the light and dark periods were maintained to the entrained day-night cycle experienced by the larvae prior to the assay. Intermittent light-dark switches and vibrational stimuli were administered to challenge the animals and record their behaviours to disturbance. Both the WT and *chrna5* mutants maintained normal diurnal circadian rhythms, exhibiting lowest activity at night (Figure 6(**f**) Cliff’s Delta = 0.514, 95%CI [0.284,0.690], *p*<0.001, Figure 6(**g**) Cliff’s Delta = 0.840, 95%CI [0.699,0.922], *p*<0.001) and highest in the morning (Figure 6(**f**) Cliff’s Delta = - 0.797, 95%CI [-0.894,-0.635], *p*<0.001), Figure 6(**g**) Cliff’s Delta = -0.703, 95%CI [-0.834, -0.514], *p*<0.001. However, the mutant fish displayed lower overall activity during each time period, with the greatest difference between the genotypes occurring during the night (Figure 6(**h**)). Conversely, during the nocturnal vibration tests, *chrna5* mutants exhibited greater motility, suggesting increased sensitivity to physical stimuli (Figure 6(**i**)). During the light-dark transition period of the assay, both genotypes exhibited a locomotor response to both the “dark flash” (light to dark), and the “light flash”, that is dark to light transition. However, the WT fish response peaked considerably higher than the mutants when the light was turned off, and maintained greater activity until the light was switched back on (Figure 6 (**j-k**)). The WT also showed a stronger response to a dark-light transition, but in this case, exhibited reduced locomotion following the light being switched on (Figure 6 (**j, l**)).

The link between nicotine consumption and feeding behaviour is well documented, with nicotine administration being associated with reduced body weight and a decrease in consumption of calorie dense foods [68,69]. Nicotine is thought to both alter the balance of orexigenic and anorexigenic peptides to change homeostatic feeding, while also exerting influence over dopamine release which modulates hedonistic feeding behaviours [70]. In addition, the CHRNA5-CHRNA3-CHRNB4 gene cluster is expressed in the arcuate nucleus of the hypothalamus (ARC) which can impact appetite directly [71,72]. Global Chrna5 receptor knockout in rodents changes preference for some food rewards, but not all [73]. To evaluate this further, we examined consumption of three types of food: a fat rich chicken egg yolk, a customised protein rich feed, and live *Paramecium caudatum*, a natural prey item of larval zebrafish in an appetite assay [38]. The assay quantified feeding by fluorescent labelling of the feed and subsequent quantification of the fluorescent signal from the gut following the feeding period (Figure 6(**m**)). Swarm Plots of the quantity of feed consumed by zebrafish across 10 repetitions of the assay for each food type revealed that the *chrna5* mutants tended to eat less fat-rich egg yolk than the WT (Figure 6(**n**) Cliff’s Delta = -0.203, 95%CI [-0.109, -0.296], *p* <0.001), but, more protein (Figure 6(**o**) Cliff’s delta = 0.312, 95%CI [0.218, 0.404], *p* < 0.001) and *Paramecium* (Figure 6(**p**) Cliff’s delta = 0.18, 95%CI [0.088, 0.274], *p* = < 0.001). Therefore, similar to the observations in rodents, *chrna5* mutants show an increased appetite for palatable food reward.

Overall, the behaviour of *chrna5* mutants differed from the WT in some phenotypes associated with human disorders often comorbid with substance dependence, such as appetite and circadian regulation.

## Discussion

Here, we generated a global *chrna5* zebrafish mutant to study the neurobehavioural consequences of dysfunction in a genetic factor associated with several frequently comorbid human disorders. *chrna5* mutants exhibited heightened tolerance to acute nicotine self-administration, a phenotype also observed in loss-of-function rodents [18,21]. This suggests that dysfunctional *chrna5* impacts tolerance to nicotine across vertebrates, and may similarly affect humans with reduced *CHRNA5* function. Notably, Genome Wide Association Studies (GWAS) have independently linked polymorphisms in the *CHRNA5– CHRNA3-CHRNB4* gene cluster to alcohol dependence [22]. Rodent studies exploring this association have yielded mixed results, with transgenic rodents expressing human *CHRNA5* polymorphism exhibiting behaviors similar to human phenotypes [26,74], but *Chrna5* knockout mice show an impact on alcohol consumption only at certain concetrations [24,25]. In contrast, zebrafish *chrna5* mutants showed an increased tolerance to both nicotine and alcohol self-administration. Therefore, the natural phenotype of zebrafish mutants aligns more closely with human GWAS.

Development of substance dependence is a multigenic, multiphasic, and multi-level phenomenon. Although our experiments of multiday nicotine or alcohol pre-treatments are relatively short, our investigation of the midbrain transcriptomics alongside acute self-administration in juvenile fish is particularly relevant in the context of Adolescent Brain Cognitive Development studies (ABCD [9,10,75]). Chronic nicotine exposure over weeks is known to desensitise nicotinic acetylcholine receptors [29,76], and influence reinforcement behaviours mediated by dopaminergic circuits [77]. Most rodent studies report that such exposure reduces inhibitory control and drug aversion, resulting in higher nicotine intake even after only a brief preexposure [28,78]. However, social suppression in nicotine self-administration has also been observed [79], and environmental enrichment and social housing conditions can reduce alcohol self-administration [80,81]. Thus, examining how genetic factors modulate these behaviours in the zebrafish - a shoaling, social species offers valuable comparative insights.

Our data showed that juvenile WT fish exhibited broad transcriptomic changes in both excitatory (glutamatergic) and inhibitory (GABAergic) neurotransmitter receptor systems following nicotine pre-treatment (Figure 4 (**j-l**)). These changes likely maintain equilibrium across reward and aversion circuits, as indicated by the nicotine self-administration profiles of pre-treated and untreated groups (Figure 4(**a**)). The transcriptomic changes seen in the WT parallel findings in adult rodents where modulation of dopamine receptors drd1 and drd3 [82,83], or NMDA receptors (grin) grin1-3 [84,85]) reduces intravenous nicotine self-administration. Unlike our expectations, however, nAChR gene expression changes were minimal (Figure 4(**i**)) with upregulation of α4 as the only exception, which may influence the dopaminergic neural function [86]. In contrast, the differential gene expression profile of the *chrna5* mutants changed minimally after nicotine pre-exposure, while their behaviour shifted towards aversion (Figure 4). These results suggest that reward and aversion to nicotine may be finely tuned by the same genetic players across vertebrates.

Unlike nicotine pre-treatment, alcohol pre-treatment induced large changes in cholinergic receptor expression in the WT fish, with small or no change in dopaminergic, GABAergic, and glutamatergic receptor gene expression. Changes were seen in non-neural genes (*chrna1, chrne, chrnd*, and *chrng [87,88]*) as well as neuronal *chrna6* and *chrna9*, both associated with nicotine [89,90] and alcohol [87,91] dependence in humans. The upregulation of GABRA6 in WT fish was interesting to note, as human variants in the gene are known to reduce sedation, though the effects on acute self-administration are unknown [92,93]. Once again, *chrna5* mutants showed a limited gene expression profile change in comparison to the WT, and showed the strongest behavioural shift, to nicotine self-administration (Figure 5 (**d-f**)). In addition, increased variance exhibited by *chrna5* mutants (Figure 5(**c**)) suggested influence over inter-individual response variability. Thus, alcohol pre-treatment induced compensatory gene expression changes in WT that were absent in the mutants. Together, these observations suggest an inverse relationship between transcriptomic changes and behavioural responses after preexposure to psychoactive substances like nicotine and alcohol for short periods. Whether longer term exposure maintains these differences and direction of change is a topic of further research. If the WT transcriptome reflects homeostatic changes, compensating for circuit functionality, then mutants lacked this protective response. If this mechanism is conserved across vertebrates, *CHRNA5* variants that reduce the function of CHRNA5 in humans could suppress compensatory biochemical changes, amplifying the direct impacts of nicotine or alcohol.

Nicotine and alcohol comorbidity is well documented in humans [50,51], and at a molecular level in animals [94], making the effects of cross treatments particularly interesting. While rodent studies commonly report that drug pre-exposure increases subsequent intake [28,49] we only observed this for nicotine pre-treatment followed by alcohol self-administration. Nicotine pre-treatment shifted both genotypes towards alcohol attraction, suggesting that the transcriptional changes induced by one substance can influence tolerance or another (Figure 4 (**d, e**)). In addition to technical differences between species, these findings may indicate the impact on the developing brains of juvenile fish. Furthermore, many genes exhibiting altered expression in the WT were linked to addiction, withdrawal, and relapse, highlighting the bidirectional nature. These included D2 and D4 receptors, which exert influence on withdrawal symptoms, place preference, and relapse to nicotine seeking behaviour but not acute nicotine response [95–97], and have human variants associated with nicotine and alcohol use [98,99]. Interestingly, D4 was also differentially expressed prior to pre-treatments (Figure S8, S9), while D3 (drd3) known to affect alcohol consumption in rodents [100], was one of the few genes with expression altered to a similar degree in both genotypes by nicotine pre-treatment. Additionally, GABRA2 and GABRA4 have several nicotine dependence associated SNPs [101], while GABRR1 and GABRR2 SNPs have been associated with alcohol dependence [102]. Finally, *GRIN2A* is associated with heroin addiction in GWAS [103]. These observations suggest that zebrafish can provide new insights into understanding the initial phases of substance dependence in young brains.

Mutations in *CHRNA5* are associated with vulnerability to anxiety, albeit only under certain conditions [57,74]. Mutant zebrafish displayed no altered anxiety phenotype in the light/dark assay (Figure 6 (**b-d**)), thus more closely matching human GWAS studies than rodent studies. CHRNA5’s role in nicotine use disorders may thus develop independently of an anxiety phenotype. Increased tolerance, in turn, could influence an increased rate of smoking. As anxiety phenotypes can vary in their manifestations and differ between social and asocial settings, additional studies examining other anxiogenic conditions will be needed to further evaluate if the dissociation manifests in other contexts.

The link between nicotine consumption and circadian rhythms is also well known [63–66]. More recent investigations are beginning to uncover the bidirectional influence of genetic variation on both [66]. The diurnal activity cycle of the *chrna5* fish mutants was similar to the WT, even if their activity was lower at various stages within it, especially at night (Figure 6 (**f-h**)). This suggests a potential role for *chrna5* in sleep quality, independently of nicotine or alcohol consumption. Additionally, the visuomotor response (VMR) to dark flashes and dark-light transitions [104] was weaker in *chrna5* mutants. As low levels of expression could be seen in the preoptic, hypothalamic, and pineal gland circuits that are proposed to influence these responses [105,106], it implies that *chrna5* may have a direct role. It is also possible that *chrna5* function is more relevant in the distributed learning circuits associated with dark flash habituation [107].

Finally, nAChR receptors have been previously implicated in the control of feeding and appetite [71,72]. Our results suggest that *chrna5* exerts some influence over appetite, independent of nicotine administration, as evidenced by the altered consumption by mutants (Figure 6 (**n-p**)). However, the direction of change was dependent on food type - mutants consumed less egg yolk, but more of a protein-rich diet, and *Paramecium caudata*. Among these, paramecia require prey hunting behaviour and coordinated locomotion making it more challenging compared to passive feed like protein-rich and egg yolk powder [108]. Notably, reduced locomotion seen in the circadian rhythm assay (Figure 6(**h**)) did not impact the *chrna5* mutants to outperform WT larvae in hunting. As the effect sizes were small in these assays, *chrna5* may contribute only minimally to appetite regulation. In rodents, *Chrna5* was amongst the lowest abundance nAChR subunits in the hypothalamic arcuate nucleus, and did not form functional receptors with β4, a known mediator of nicotine induced appetite reduction [109]. Since these feed types differ dramatically in form, nutritional quality, and motility further studies are necessary to understand what among macronutrient content, visual cues, or hunting drives mutant behaviour [110].

## Conclusion

This whole-body *chrna5* mutant zebrafish exhibits aligned phenotypes with human conditions, making it a valuable and practical model for investigating the neurobiological basis of development of substance use disorders (SUD). Short term pre-exposure to multiple substances revealed novel cross-substance effects on nicotine and alcohol self-administration in these mutants, highlighting the impact of prior substance exposure on subsequent drug use behavior. Parallel transcriptomics showed altered expression of dependence associated genes in wild type fish that were absent in mutants, suggesting a disruption to homeostatic responses. Together, our results suggest that disruption of *chrna5* increases susceptibility to substance dependence, impairs adaptive responses, and affects behaviours such as appetite and circadian rhythms, which are frequently altered in humans in the context of SUD.

## Materials & Methods

### Fish husbandry

Zebrafish (*D. rerio*, ABWT) were housed, bred, and reared at the ZebraFish Facility (Institute of Molecular and Cell Biology, A*STAR) in groups of 20–30 in 3 L tanks at 28°C. Fish were kept under a 14:10 light dark cycle and fed as per standard operating procedures of ZFF. All experimental protocols involving zebrafish were approved by the Institutional Animal Care and Use Committee (IACUC) of the Biological Resource Center at A*STAR. Approved experimental protocols (IACUC #201529, #231808, #231797) were followed.

### Generation of chrna5 mutant zebrafish

Zebrafish gene editing by CRISPR-Cas9 is comprehensively described by Hwang *et al. [111]*. In brief, CRISPR targets for *chrna5* were determined using the web tool ‘chop-chop’ for the exon 8, targeting between the 3rd and 4th transmembrane helices to disrupt key functional domains. Primers for generating sgRNA are detailed in Table 1. Zebrafish embryos (nacre background) were injected at the 1-cell stage with 1 nL of a mixture containing 1 μL sgRNA (≈ 4-5 μg) and 1 μL Cas9 mRNA (≈1 μg). Fish were genotyped by fin-clipping and sequencing of genomic DNA at 10 weeks post-injection to identify founder fish (F0). The identified founder fish were outcrossed to AB background WT fish to obtain F1 embryos. F1 embryos from each outcrossed family were collected and some of the embryos were genotyped. Once germline transmission was confirmed, the remaining embryos were grown to adulthood. Homozygous *chrna5* mutants from the F3 generation onwards were used for all subsequent experiments, henceforth referred to as *chrna5*^*-/-*^.

**Table 1.**
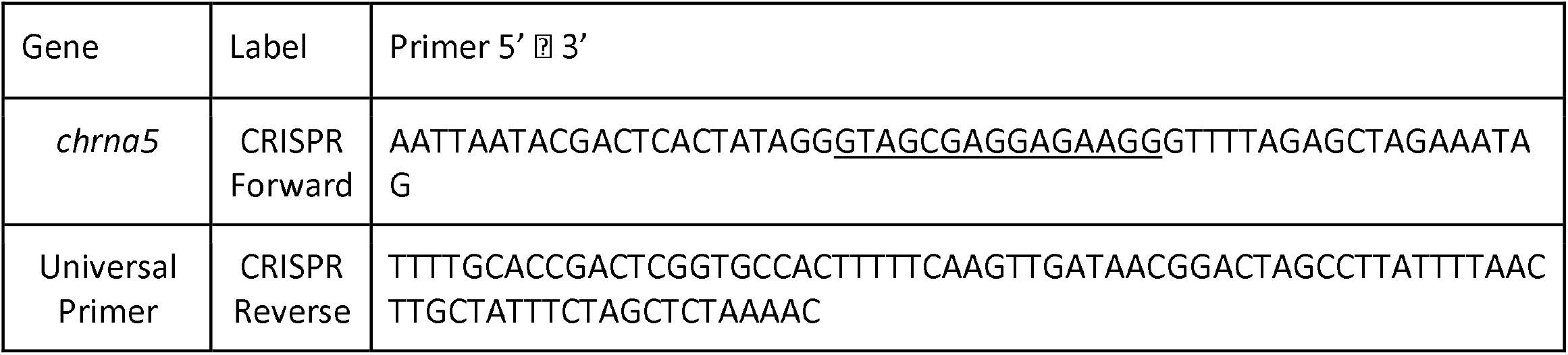
Primers for single-guide RNA template. CRISPR targets are underlined

### Self-administration for zebrafish assay (SAZA) apparatus setup

Full descriptions of the tanks and apparatus used for the SAZA assay can be found in [40] and [41]. In brief, the assay tanks were constructed of 3mm thick opaque acrylic (dimensions: 35 × 75 × 30mm, width × length × height). One narrow end of the tank was divided by a sheet of the same material, 30mm in length, to split that end into two halves, while still permitting access by the fish (Figure 2(**a**)). The tank was placed on top of an LED lightbox, providing 5000 lux at maximum intensity (LightPad LX Series, Artograph, USA). Videos were captured at 30 frames per second using an acA2040-90μM USB3.0 Basler camera. Nicotine, alcohol and control solutions were dispensed by gravity from 10ml syringes, mounted vertically an equal distance above the tank, through silicon tubing (outer diameter: 1/16”, inner diameter 1/32”) into the corner of their designated zone. Dispensing was mediated via solenoid pinch valves, set to open only when a tracked fish entered the designated zone (Automate Scientific, USA, SKU: 02-pp-04i) (Figure 2(**a**)). Fresh system water was consistently supplied to the tank via additional silicon tubing at the opposite end of the tank to the stimulus/control delivery zones. Water was also removed from the tank at a matching rate (∼2ml per minute) by gravitational siphoning, with the outflow tubing positioned at the end of the dividing sheet, outside the stimulus/control delivery zones (Figure 2(**a**)). This created a constant gentle flow of water through the tank, while also preventing the dispensed solutions from escaping their designated zones (Figure 2(**a**)). Custom LABVIEW software, CRITTA (http://www.critta.org), was used to track fish movements as described by Krishnan *et al*. (2014) [112].

### SAZA

The SAZA assay was performed as described by [40] and [41]. Each tank was filled with 40 ml of system water, with the system water inflow and outflow tubes, and the stimulus and control dispensing tubes, inserted into their respective positions. Juvenile zebrafish (30-35 dpf), naive to SAZA, were taken from the husbandry tanks, and one was added to each SAZA tank while minimising disturbance. Each fish was given approximately five minutes to adjust and recover post-transfer, until consistent swimming behaviour had resumed and the fish had explored all zones of the SAZA tank. Fish that did not exhibit consistent swimming behaviour after the recovery period were removed from the tank and replaced with a new fish. The recording and tracking scheme was then started, lasting for 24 minutes and consisting of three consecutive periods; three minutes of pre-exposure, 18 minutes of self-administration stimulus delivery, and three minutes post-exposure. Only during the 18-minute stimulus delivery period would entry to a designated stimulus/control zone trigger dispensing of the corresponding solution (Figure 2(**a**)). The stimuli delivered were 0/5/10/20% of absolute alcohol, or 0/10/500μM nicotine hydrogen tartrate salt (cat #6019-06-3 Sigma) diluted in system water. These source concentrations were diluted in the stimulus zone, with the concentration increasing over time with repeated fish entries, estimated from ∼40x to 10x diluton with nicotne, and measured at ∼70x to ∼30x with alcohol. This resulted in mean stmulus chamber concentratons during SAZA of 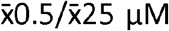 nicotne and 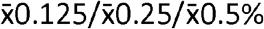 alcohol, respectively, as outlined in Table S2. These terms were used for the remainder of the work. Following each assay, the tank was emptied of all liquid, rinsed, and refilled before adding the next fish to prevent contamination of the subsequent assay. The volume of each solution dispensed by each fish was recorded. Periodically throughout the experiment, the zone designated to deliver the stimulus solution was randomly assigned to be either the left or right, and changed such that ∼50% of the fish were tested in each configuration in order to reduce bias. Approximately 30 fish were assayed per concentration of alcohol or nicotine. Following each assay, the fish were euthanized. In some cases, zebrafish were pre-treated with exposure to alcohol or nicotine solutions before being assayed. For these assays, each fish was immersed in either 1µM nicotine or 1% alcohol, diluted in system water, for one hour for seven consecutive days. Following this pre-treatment, these fish were returned back to the facility for usual husbandry. On the eighth day, the fish were subjected to SAZA, either using the same substance as introduced during pre-treatment, or the opposite, creating four total conditions (Figure 3(**a**)).

### SAZA Data analysis

CRITTA tracking data from SAZA were processed by custom Python scripts. These provided data on time spent in each zone, mean velocity in each zone, number of entries, and mean time per entry used in these analyses. The data were separated into three major time groups; pre-exposure, total stimulus delivery period, and post-exposure. The stimulus delivery period was also subdivided into three minute segments (0-3m/3-6m/6-9m, and so forth) for some further analyses. The preference index was calculated for the total stimulus delivery period, based on the volume of each solution dispensed, by the following formula; PI = (VolS - VolC) / (VolS + VolC), where VolS = volume dispensed in stimulus zone (ml) and VolC = volume dispensed in control zone (s). This gives the value for PI a maximum range of + or - 1 indicating more volume dispensed, relative to the total, in the stimulus or control, respectively. For example, a PI of +1 would indicate 100% of the total volume dispensed was in the stimulus zone during the assay, while a PI of 0 would indicate equal volumes were dispensed in each zone.

### RNA extraction

Six-month old adult zebrafish were euthanized, and the brains were dissected in 1x PBS pH 7.0, selecting the forebrain and midbrain regions (excluding the olfactory bulb, telencephalon, and hindbrain regions). Brain tissue was homogenised using a micro tube homogenizer (Thomas Scientific) in a Trizol lysis buffer (ThermoFisher #15596026). RNA was then extracted using PureLink® Micro Kit (ThermoFisher #K310250) according to the manufacturer’s protocol. The concentration and purity of the extracted RNA were determined by NanoDrop™2000 Spectrophotometer (ThermoFisher). Each brain sample yielded around 100 ng/μL RNA with a ratio of absorbance reading 1.9-2.0 at 260/280 nm. Some RNA samples were sent for bulk sequencing, detailed below. The remaining purified RNA was reverse transcribed by SuperScript II First strand Synthesis System (Invitrogen #18091050) using 100 ng/μL of purified RNA to obtain approximately 1000 ng/μL of first strand cDNA. Negative controls containing no reverse transcriptase were set up for each sample to check for genomic DNA contamination. The cDNA was diluted with nuclease-free water to 100 ng/μL and used for qRT-PCR.

### RNAseq

The quality of each RNA sample was verified before sequencing by gel electrophoresis and bio-analyser. Samples that passed QC were sent for sequencing by BGI. An mRNA cDNA library made up of paired-end sequencing reads of 150 bp in length was generated on the Illumina HiSeq instrument. Raw reads were first processed to remove adapters, poly-N sequences, and reads with low quality from the raw data. All downstream analyses were based on the clean data with a Phred score of 39, indicating a 99.9% base call accuracy. Further analysis was performed in Partek™ Flow™ Explore Spatial Multiomics Data using Partek™ Flow™ software, v11.0. Reads were aligned to GRCz11 genome assembly by STAR aligner, default parameters, and subsequently filtered to a minimum mapping quality score of > 30. Gene counts were normalised by median ratio, followed by differential expression analysis in DeSeq2. Significance thresholds were set as follows: up/down-regulated genes = p < 0.05 & fold change > ±2. Provisional up/down-regulated genes = p < 0.05 & fold change < ±2. Non-significant genes = p > 0.05. Custom gene lists and sets for analysis can be found in Table S10. Gene set enrichment analysis (GSEA) was performed using the GRCz11-GO database, 100 permutations. Significance thresholds were set at p < 0.05 and FDR < 0.25.

### qRT-PCR

The qRT-PCR amplification mixtures (20 μL) contained 100 ng of cDNA, 10 μL 2x GoTaq®qPCR Master Mix (Promega #TM318), and 300 nM forward and reverse primers. The primers used were designed using Primer3 (v.0.4.0) and are detailed in Table 2.

**Table 2.**
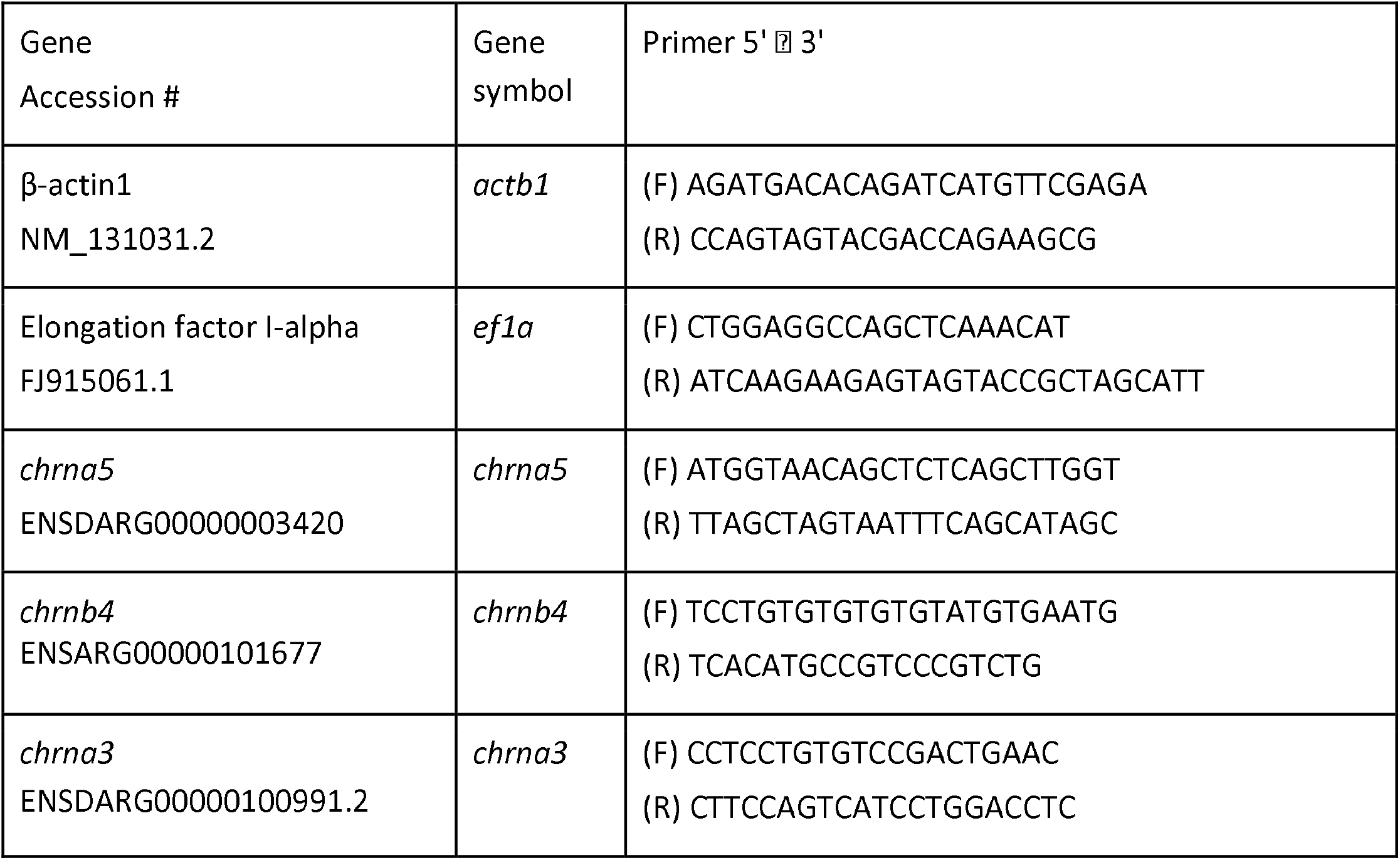
Primers used for qPCR of *chrna5*, and housekeeping genes.

Reactions were performed using an Applied Biosystems 7500 Real-Time PCR system, in 96-well plate format. The cycling conditions were as follows: 95°C for 10m to activate, then 40 cycles of 95°C for 15s, 60°C for 30s. After completion, a melt curve was run at 65-95°C for 5 seconds per step. All PCR efficiencies were between 90-110%. Primer specificity was validated by a single peak on the post-PCR melt curve and single-band after electrophoresis. The relative gene expression levels were normalised to the housekeeping genes and analysed using the adjusted delta-delta CT method, described by Hellemans and Mortier et al. (2007). All data were analysed as described below.

### Statistical analyses

Data in this study were analysed and presented following the principles of hybrid effect size plus *p* value [113,114]. Briefly, a P value as a measure of significance is improved upon by the supplementary reporting of effect sizes, means, and 95% confidence intervals [114]. The assumptions of homoscedasticity and normality typically recommended for parametric testing were not met consistently for the SAZA data. As such, unpaired data were compared by non-parametric Cliff’s delta effect size and two-sided permutation T-tests (5000 reshuffles) with a significance level of 0.05. Repeated measures data were compared by mean difference and paired permutation tests under the same parameters. Vibration assay longitudinal data were analysed using a linear mixed effects model, with pixels travelled as the response variable, genotype and vibration intensity as fixed effects, and individual ID as a random effect to account for repeated measures. Analyses were performed in R using the ‘dabestr’ (v2024.12.24) and ‘lmerTest’ (v3.1-3) packages. If multiple comparisons were made, family wise error was corrected for by applying the Holm-Bonferroni step down method to *p*-values. Outputs of these statistical analyses can be found in supplementary tables. Where data were presented as Gardner-Altman and Cummings estimation plots, the results are reported as Cliff’s delta, or Paired mean difference = X, 95% CI [lower, upper], *p* = Y. For Cliff’s delta, effect sizes greater than ± 0.4 and a *p*-value of < 0.01 were considered meaningful and practically relevant in this study. Smaller Cliff’s delta effect sizes of ± 0.2 to 0.4 with accompanying *p*-value of < 0.05 were considered provisionally meaningful, pending reproduction or further investigation [115,116].

### Hybridisation chain reaction, RNA-fluorescent in-situ hybridisation (HCR™ RNA-FISH)

Larval zebrafish for HCR™ RNA-FISH 3.0 (HCR) [117] were collected as follows. During development, 300µL of 0.3% PTU (Phenylthiocarbamide, Sigma) was added to each petri dish containing larvae every alternative day from 3 dpf until 14 dpf to reduce pigmentation. Following this, larvae were quickly rinsed with 1x PBST, euthanized and fixed in 4% formaldehyde, diluted in 1X PBST, at room temperature for 60 minutes with shaking. Once fixed, the larvae were washed with 1X PBST for three minutes, three times. Larvae were then permeabilized using a detergent solution (1% SDS, 0.5% Tween 20, 150mM NaCl, 1mM EDTA, 0.05M Tris-HCL, pH 7.5) at 37°C for 30 minutes and washed for three minutes, three times each with 1X PBST and then 5X SSCT. Finally, the larvae were transferred to 30% probe hybridization buffer (30% formamide, 9mM citric acid, 0.1% Tween 20, 50µg/mL heparin, 1x Denhardt’s solution, 5X SSC) and incubated at 37°C for 30 minutes. At this stage, the larvae were stored at 4 degrees for up to one month in the dark or used directly for HCR.

HCR probes against *chrna5, neurod1*, and *nrp1a* were prepared using an in-house Excel-based program and synthesized by IDT (see Table S1 for details). Probes were prepared in a cocktail containing 30ul (10 µM) of each probe set against each gene in 1000 µl of 30% probe hybridization buffer. The in-situ hybridization experiments were carried out in an automated robotic fluidics system. The larvae were transferred to the automated fluidics robot with the primary, secondary, and wash buffers, and a script to perform the reaction was loaded (preprogrammed in the system). The larvae were first incubated with the probes at 37°C for 8 hrs, then washed five times with 30% probe wash buffer (30% formamide, 9mM citric acid, 0.1% Tween 20, 50µg/mL heparin, 5x SSC) at 37°C and then 5X SSCT twice. For the secondary reaction, HCR hairpins diluted and prepared in an amplification buffer (5X SSC, 0.1% Tween 20, 2.5% dextran sulphate) were added and incubated for 3 hours in the dark in the automated system. The larvae were then washed with 5X SSCT, eight times, for 2 mins each. After the reaction, the samples were removed from the chamber and mounted with the cover slip on the dorsal side in 70% glycerol-based mounting media and imaged on an Olympus FV3000 confocal microscope with the settings consistent between fish. Images were post-processed first on Olympus viewer FLUOVIEW FV31S-SW software with linear intensity adjustments of 150 - 4095 for all genes. Images presented for *nrp1a, neurod1* and *chrna5* had no further processing beyond cropping to the ROI and addition of scale bars.

### Protein extraction

Protein was extracted from the brains of adult fish, aged 6 months. The fish were euthanized, then subsequently dissected in 1x PBS pH 7.0. Brain tissue was transferred to 400 µl of cold, 6 M urea buffer with added protease inhibitors, and homogenized until the solution was clear of visible debris. The protein samples were then briefly vortexed and centrifuged for 30 seconds to pellet debris. The samples were then sonicated with three 30 second pulses at one minute intervals using a probe sonicator on ice. The samples were then incubated on ice for 20 minutes before centrifuging at 14,000 rpm for 20 minutes at 4 °C. The supernatant was then transferred to a new tube, aliquoted, and stored in -80°C until further use. Protein concentrations were measured using the Pierce BCA Protein assay kit (Thermo Fisher Scientific), following the manufacturer’s protocol in 96 well plate format. BSA standards were serially diluted in 6M urea buffer from a working concentration of 2000 µg/ml to 25 µg/ml to create a standard curve. In each well, 25 µl of diluted BSA standard or protein sample was added, and each topped up by 200µl of BCA working solution. The plate was covered and incubated for 37°C for 30 minutes, before absorbance at 562 nm was determined by a plate reader. From one round of protein extraction, approximately 0.8 mg/ml was extracted.

### Western blots

Western blots were performed on extracted brain proteins by Wes system (ProteinSimple, product number 004-600) using a 12-230 kDa Separation Module 13-capillary cartridge SM-W002. Samples were diluted to 1 µg/µL in 6 M urea buffer, mixed in fluorescent 5x Master Mix, including DTT, and denatured at 95°C for five minutes. The samples, primary antibodies (1:1000 diluted in Wes antibody diluent), HRP-conjugated secondary antibody (1:4000 diluted in Wes antibody diluent), and chemiluminescent substrate (luminol-S and peroxide) were added onto the plate, in addition to the biotinylated ladder, antibody diluent and wash buffer included on the plate by the manufacturer. Default instrument settings were used, stacking and separation at 475V for 30 minutes, blocking 5 minutes, primary antibody incubation, 60 minutes, and secondary antibody incubation, 30 minutes. The chemiluminescence detection was set for 30 minutes total, with imaging at 2-minute intervals. The electropherograms were first checked to see if manual correction was needed for peak detection, and then the band intensity results were exported. The following primary antibodies were used: for the reference protein, monoclonal anti-Tubulin antibody, beta, clone KMX-1 (MAB3408) Sigma, produced in mice. For *chrna5*, Monoclonal anti-CHRNA5 antibody (AV34967-100UG) Sigma, produced in rabbit, Table 2.3.3.3. The secondary antibodies used were polyclonal anti-mouse goat antibody (P0447 DAKO) HRP conjugated and polyclonal Anti-rabbit (sc-2004 Santa Cruz Biotechnology), produced in goat HRP Conjugated.

### Light/Dark preference assay

The light/dark preference assay, designed to test nyctophobia/photophila as a proxy for anxiety, was set up and performed as described by [118] and [119]. Four transparent rectangular plastic boxes (70mm x 40mm x 15mm, L x W x H) were each filled with 50ml of system water. The boxes were placed inside a cabinet, covered with a cloth to prevent any light interference. Each box was divided into light and dark sides, each 35 mm in length, by using an iPad (Apple 7^th^ generation, display at highest light intensity) underneath to project an equal light/dark split image (Figure 6(**a**)). A camera (ACa2040-90 uM USB 3.0: Basler resolution 800 × 600 pixels) with infrared filter was placed overhead, which recorded the movements of 12-14 dpf larvae. To prevent social interference, dividers were placed in between the four chambers to isolate the fish completely. The setup was further illuminated by placing two infrared LED bars on either side of the tanks (Figure 6). The camera output was recorded using Pylon viewer Ver 6.1.1. All trials were conducted between 9am-6pm. Prior to the assay, larvae were either not treated with any stimulus. The larvae were quickly and gently added to the illuminated rectangular boxes, with one larva per box, four boxes at a time. A total of ∼36 individuals per treatment and genotype were assayed. Larvae were allowed to acclimate in the chamber for ten minutes until free, uninhibited swimming was observed before starting the video recording. Then, the larvae were filmed for ten minutes in complete light, followed by a further ten minutes in the light and dark setting (Figure 6(**a**)). Each fish was tracked in real time by an in-house custom-written Python code using the opencv library [118]. Each recording lasted 10,000 frames at 16 fps, totalling approximately 10 minutes. The x-y coordinates of each larva in each frame were exported. Automated analyses [118] were applied to all larvae, however, individuals that moved less than 50 mm during the total assay duration were excluded from the final data.

### Circadian Rhythm assay

The circadian assay was performed in 48-well plates, with each well containing 1ml of system water. These plates were placed inside an opaque rectangular box to avoid outside light interference, on top of an illuminated light box, surrounded by four IR LED bars (LBS2-00-080-2-IR850-24V, 850nm: TMS Lite). A camera (Aca2040-90µM USB 3.0: Basler) with an IR filter was positioned above the plate to track the movement of each 7-10 dpf larva. At the onset of recording, ∼15:00-15:30 pm, the larvae were kept in continuous ambient light. The illumination was switched off at 22:30 pm, and switched on again at 08:30 am the next day, and the assay ended after 24 hours. Larval movements were tracked by an in-house custom-written script in Python 2.7 [120], incorporating functions from the OpenCV library to control the Arduino microcontroller and video track larvae movement. Videos were captured at 576 × 854 px at 10 fps, and the background subtraction method was applied to obtain the X and Y coordinates of the fish. The protocol is further described in detail in [120]. The following key metrics were quantified: total seconds spent moving per minute, mean velocity, mean resting period, and average distance moved per fish. Activity thresholds were defined as such: an active bout was indicated when the larvae moved five seconds or more per minute. Meanwhile, a resting, or inactive bout, was indicated when larvae moved four seconds or less for one minute. For the assay duration, larvae were either not treated with any stimulus. Two other behavioural assays were also conducted during the circadian assay duration, testing the startle responses to light/dark, and to vibration. The light/dark startle response test occurred for five, 30 minute light/dark cycles for a total of five hours, from ∼17.30 – 22.30 pm (Figure 6(**j-l**)). The vibration test consisted of six, 20 minute cycles, two per hour. Within these cycles, 18 shocks were administered per hour for nine different intensities, first from 0-100% ascending, and subsequently descending, from a period of ∼1:00-7:00 am (Figure 6(**i**)). Vibrations were controlled through a speaker via an Arduino microcontroller board by a custom Python script (Python 2.7, C[120]. The highest intensity was represented by 50% on the computer audio output. The time interval between each vibration was 30 seconds, which prevented behavioural habituation. The larvae were recorded as responding if they moved more than 7 pixels (∼ 4 mm), and the average percentage of responding fish was calculated for each different intensity. The observed values were corrected against baseline locomotor activity, as further described by [120].

### Feeding Assay

Food types for the various feeding assays were prepared as follows. *Paramecium caudatum* was cultured and harvested weekly as described in the zebrafish book [121], and ‘*Paramecium* Recipes for Large and Small Facilities’. Briefly, paramecia culture supernatants were collected in a clean 50 ml falcon tube to remove any yeast accumulated at the bottom, then centrifuged (Eppendorf) at 4°C, 15 minutes, 2500 rpm to isolate the pellet. The resulting supernatant was removed and the paramecia pellet was resuspended in 1 mL of system water. Next, 2.5 µL of stock DID dye (Invitrogen) in 22.5 µL 100% ethanol was added to the resuspended paramecia and incubated for two hours at RT under nutation. Subsequently, the mixture was centrifuged at 13,000 rpm for five minutes at RT. The alcohol supernatant was removed, and the pellet resuspended in 1mL of system water under nutation for 15 minutes. The mixture was then briefly vortexed and added onto the rotator again for a further ten minutes to thoroughly mix the paramecia with system water. Finally, 1mL of this dyed paramecia solution was added to 3 mL of system water and used to feed four plates of larvae (1 mL per plate). Egg yolk from chicken (#cat E0625 Sigma) was purchased and prepared as described for the paramecia, but with the following adjustments. For egg yolk powder, 10mg was measured and mixed with 1ml of system water and 1 µL DID stock dye. The tube was vortexed and incubated for 2 hours at RT under nutation. Subsequently, the supernatant was removed, without disturbing the pellet. The pellet was then resuspended in 1 ml of system water under nutation for 15 minutes, ready for use. The protein-rich powder, custom formulated and a gift from Caroline Wee lab, was prepared in the same manner as the egg yolk powder.

To perform the assay with paramecia, the larvae were subjected to two “training days” of feeding in the assay petri dish with 1ml of paramecia (30-40 larvae per petri dish) overnight. The assay was performed at 7 dpf. Larvae were first fed with excess unlabelled paramecia for 90 minutes during the first feeding session, then washed and fasted for 120 minutes. After the fasting period, the larvae were transferred into smaller, 35mm petri dishes, each filled with 4 mL of system water, before adding 1 mL of labelled paramecia. Larvae were given 90 minutes for the second feeding session, after which they were cold anaesthetized and fixed in 4% PFA in 1x PBST overnight at 4°C. Larvae were washed three times in 1x PBST the next day before mounting and imaging (Figure 6(**m**)).

The assay was performed in a similar manner with egg yolk or protein-rich powder as the food, but with the following adjustments. On the training days at 5 and 6 dpf, the fish were instead fed with a small quantity of 150/250 micro ground dry algae (Zeitger) for 30 minutes. The algae were then washed away, and replaced with fresh system water overnight. At 7 dpf, the first feeding session lasted for only 30 minutes, before the fasting period of 2 hours. For the second feeding session, 500µl of labelled-egg yolk, or protein rich powder, was added in petri dishes for 30 minutes, before fixing. To prepare for imaging, the fixed larvae were washed three times in 1x PBS and arranged per genotype (25-30 fish) in a 35mm glass bottom dish in a circular pattern, avoiding larvae touching each other. To measure the amount of food consumed by larvae, the intestinal fluorescence signal was captured using a Leica M205 FA fluorescent stereoscope. Corresponding brightfield and fluorescent images were captured, Cy5 filter 651 nm, in a covered, dark room. The brightfield images were captured at 20 ms exposure at 40% intensity, whilst Cy5/mcherry exposure was 100ms, 100% intensity. The fluorescence intensity was measured by an in-house written code developed by Cheng et al. [120], which correlated to the total amount of food consumed by each larva. The segmentation method used for fluorescence quantification has been described in full by [120], and the code is maintained and can be found at https://github.com/CarolineWeeLab/EZgut [120]. The mean fluorescence intensity was obtained for each larva and normalized against the controls.

## Supplementary Materials

Contains 9 supplementary figures and 10 supplementary tables.

## Author Contributions

Conceptualization, A.S.M, T.G, and J.R; methodology, T.G., C.K., T.D.B, J.R. and J.W.C.; formal analysis, T.G, J.R.; investigation, T.G., J.R., and C.K., J.W.C; resources, A.S.M; writing—original draft preparation, T.G., and J.R; writing—review and editing, All.; visualisation, T.G. and J.R.; supervision, A.S.M.; project administration, A.S.M; funding acquisition, A.S.M.

## Data Availability Statement

The datasets used to generate the figures are included within the article and its additional file(s). Additional data are available from the corresponding author upon request.

## Acknowledgment

We would like to thank the zebrafish fish facility (ZFF) staff at the IMCB, A*STAR for their assistance with fish husbandry; Drs. Ruey Kuang-Cheng and Caroline Wee for assistance with conducting circadian rhythm and appetite regulation assays; Li Xinrui for generating customized Python analysis scripts.

