## Supplementary Material for "Role of *chrna5* in multi-substance preference and phenotypes comorbid with the development of substance dependence"

Goel and Raine et al., 2025. Contains 9 Supplementary figures and 10 Supplementary tables.

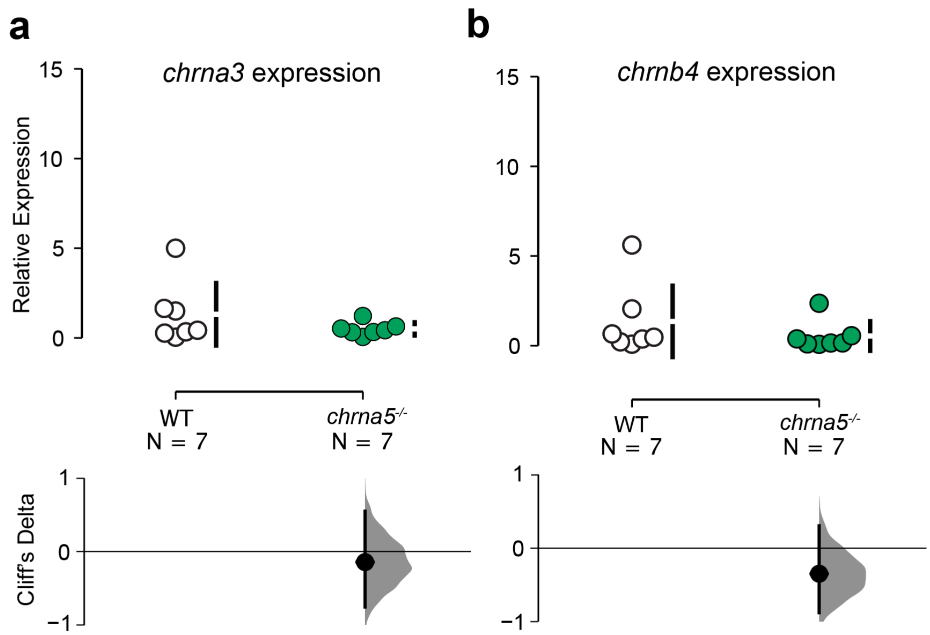

**S1. Relative gene expression for co-localized genes chrna3 and chrnb4**
A) Relative gene expression for chrna3 in WT and chrna5$^{-/-}$ mutants B) Relative gene expression for chrnb4 in WT and chrna5$^{-/-}$ mutants using gPCR analysis.

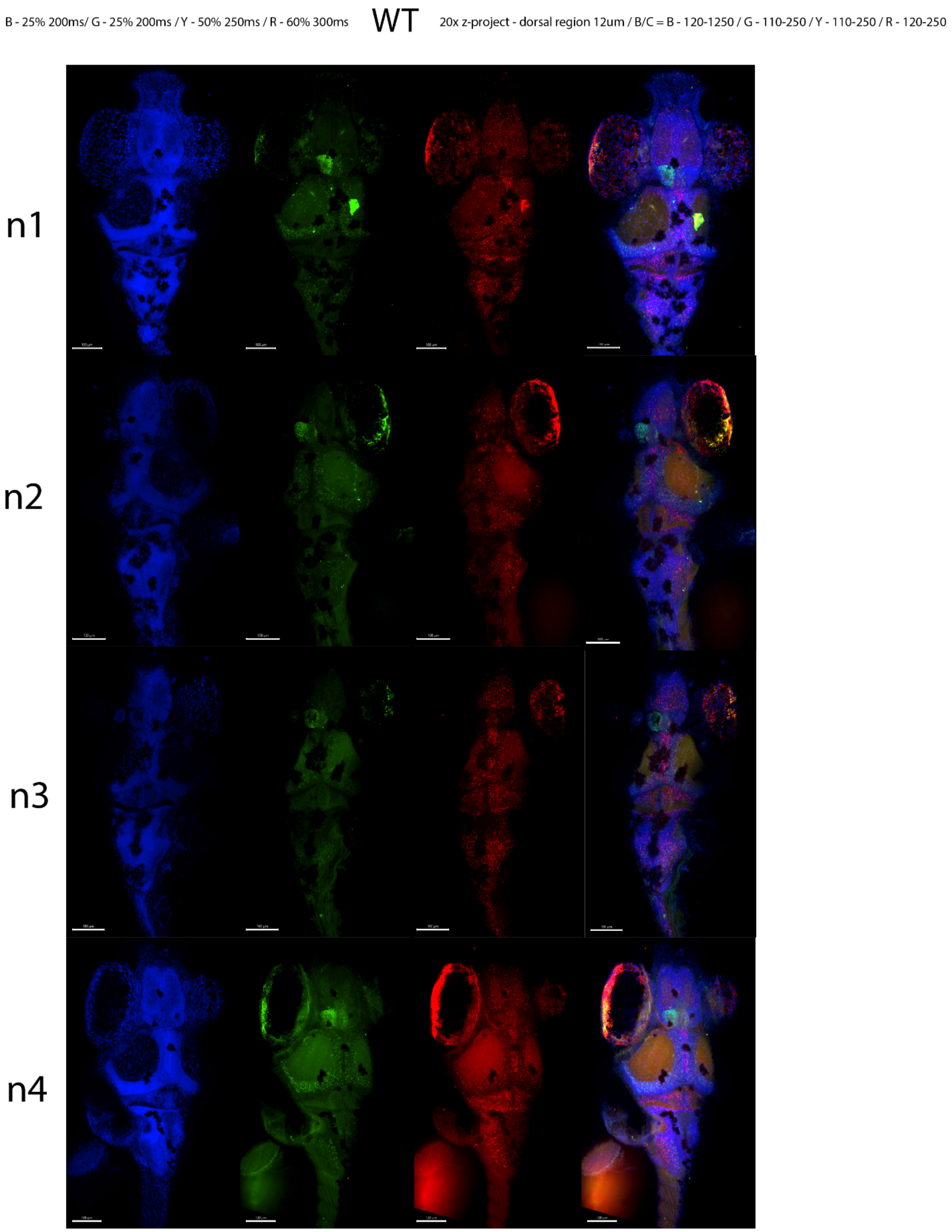

**S2. WT gene expression from HCR**
Additional HCR WT images (n=4) mounted dorsally for DAPI, nrp1a, NeuroD and chrna5. Blue = DAPI, Green = nrp1a, yellow = NeuroD and red = chrna5.

B - 25% 200ms/ G - 25% 200ms / Y - 50% 250ms / R - 60% 300ms 20x z-project - dorsal region 12um / B/C = B - 120-1250 / G - 110-250 / Y - 110-250 / R - 120-250

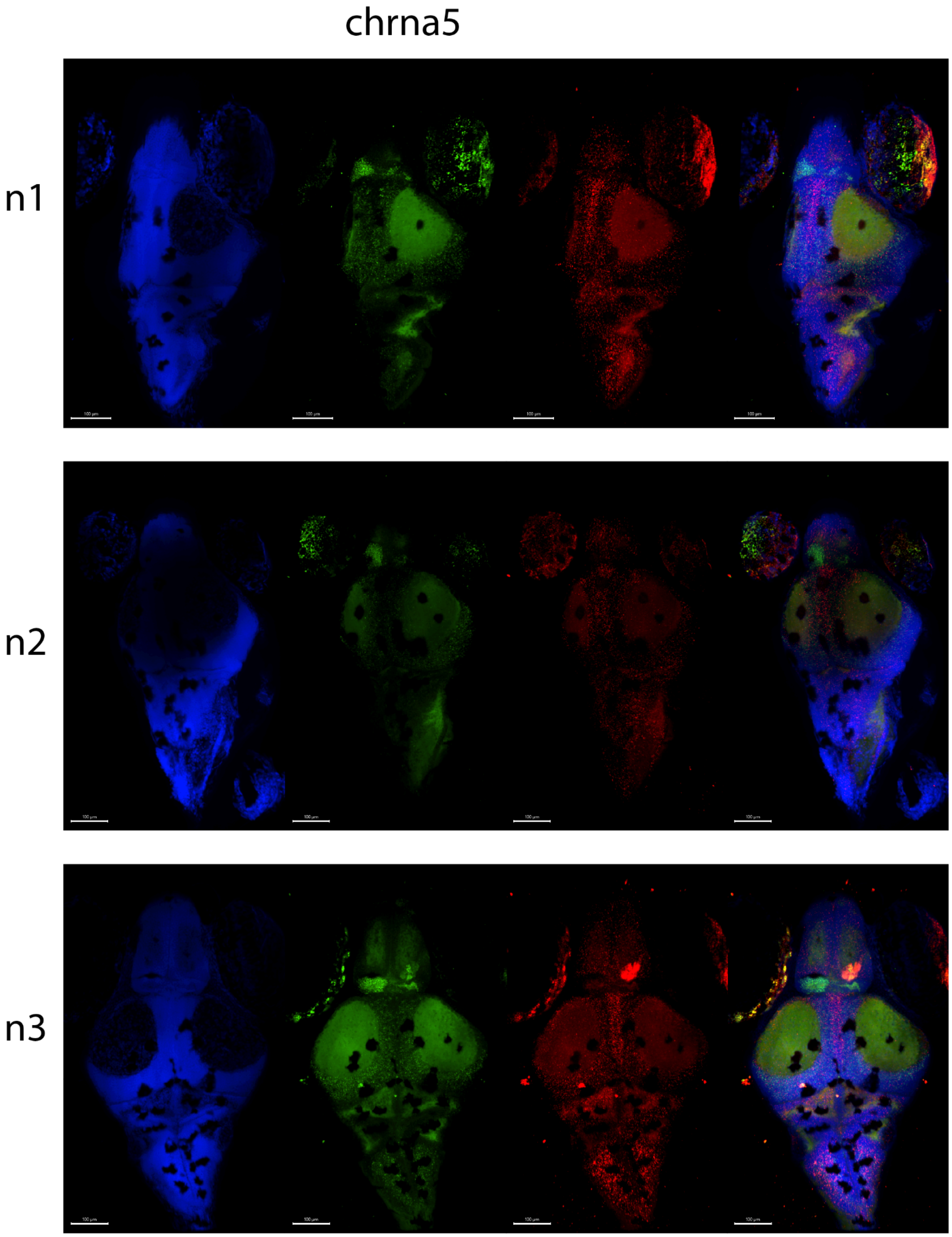

**S3. chrna5 MUT gene expression from HCR**

Additional HCR WT images (n=4) mounted dorsally for DAPI, nrp1a, and chrna5. Blue = DAPI, Green = nrp1a, red = chrna5.

**Table S1.** Probe cocktail sequences

| Probe name (gene) | Sequence |
|---|---|
| B3_P1(nrp1a)_1 | GTCCCTGCCTCTATATCTTTGTAATCACCCATATGCATTTCTGAG |
| B3_P1(nrp1a)_2 | GTCCCTGCCTCTATATCTTTGCACTCTCGGTCTTCCAGGTCAAAG |
| B3_P1(nrp1a)_3 | GTCCCTGCCTCTATATCTTTTTCCACAATATTTGCCCACCAGCTG |
| B3_P1(nrp1a)_4 | GTCCCTGCCTCTATATCTTTGTCTCGTAGTCGGACACAAACTTGA |
| B3_P1(nrp1a)_5 | GTCCCTGCCTCTATATCTTTGGTGAAGTTCCTGGAACATTCTGGA |
| B3_P1(nrp1a)_6 | GTCCCTGCCTCTATATCTTTTGAACGTGCAGTCCAAATTATTGGG |
| B3_P1(nrp1a)_7 | GTCCCTGCCTCTATATCTTTTGCGTGTCTGGCTCCAGCTCAAAAC |
| B3_P1(nrp1a)_8 | GTCCCTGCCTCTATATCTTTTGGACCAACTCCAGGGAATCCGTCC |
| B3_P2(nrp1a)_1 | AAAATCCTCTGGTTGGGTCCTGGAGTTCCACTCAACTTTAACCCG |
| B3_P2(nrp1a)_2 | GTCTCTCACTTCCACATAGTCATATTTCCACTCAACTTTAACCCG |
| B3_P2(nrp1a)_3 | ACGAGACCACCGGAGATGGAGCGATTTCCACTCAACTTTAACCCG |
| B3_P2(nrp1a)_4 | TAGCGGATGGAGAATCCGGCACCGTTTCCACTCAACTTTAACCCG |
| B3_P2(nrp1a)_5 | GGGCGACTTGATGACTCCGCTGCTGTTCCACTCAACTTTAACCCG |
| B3_P2(nrp1a)_6 | TTTCTGACATCTTAGGAGCAAAGATTTCCACTCAACTTTAACCCG |
| B3_P2(nrp1a)_7 | TATCGGCAGAAGACTCCGGCGGGCGTTCCACTCAACTTTAACCCG |
| B3_P2(nrp1a)_8 | ATTCTGTCCGCAGTATCTGCCGATGTTCCACTCAACTTTAACCCG |
| B1_P1 (chrna5)_1 | GAGGAGGGCAGCAAACGGAAGTCGTCATCAGCTGGTTCTTCTCAT |
| B1_P1 (chrna5)_2 | GAGGAGGGCAGCAAACGGAAGATGCCCAGGTAATGATCCGGATCC |
| B1_P1 (chrna5)_3 | GAGGAGGGCAGCAAACGGAATACCATCTGCATTGTCATAGAGCAC |
| B1_P1 (chrna5)_4 | GAGGAGGGCAGCAAACGGAATTGGCAGGAGGAGTCCAGGAGATTG |
| B1_P1 (chrna5)_5 | GAGGAGGGCAGCAAACGGAAGAACTTCATGGAGCAGTTTTGAAGG |
| B1_P1 (chrna5)_6 | GAGGAGGGCAGCAAACGGAAAGTCACGTTTGTCCACATGCACGTC |
| B1_P1 (chrna5)_7 | GAGGAGGGCAGCAAACGGAAAAAGTCCCATCTCTTCTCAAACCCC |
| B1_P1 (chrna5)_8 | GAGGAGGGCAGCAAACGGAAGATGAGGAAGAGTGTGTAGAAGAGA |
| B1_P2 (chrna5)_1 | GTCCATTCCTGCTTCATCCACACATTAGAAGAGTCTTCCTTTACG |
| B1_P2 (chrna5)_2 | GGAGTCTGAAGGCACCCTGATAGAGTAGAAGAGTCTTCCTTTACG |
| B1_P2 (chrna5)_3 | CTGCTTTTGTCACTGTGGCCTCGAATAGAAGAGTCTTCCTTTACG |
| B1_P2 (chrna5)_4 | ACATCAATGGTGCAGGCAGATTTATTAGAAGAGTCTTCCTTTACG |
| B1_P2 (chrna5)_5 | CTGGGAGCCATCATAGGTCCAAGAATAGAAGAGTCTTCCTTTACG |
| B1_P2 (chrna5)_6 | CGATCTCCCATTCGCCATTGTCAAATAGAAGAGTCTTCCTTTACG |
| B1_P2 (chrna5)_7 | AAGGAGTACGTGATGGATGGATAGATAGAAGAGTCTTCCTTTACG |
| B1_P2 (chrna5)_8 | CAGGAAAGACAATCCAATGCAGGGGTAGAAGAGTCTTCCTTTACG |

B1 probes used Alexa fluorophore 546, B3 probes used Alexa fluorophore 488. P1 and P2 probes are paired together of the same gene (from 1-8)

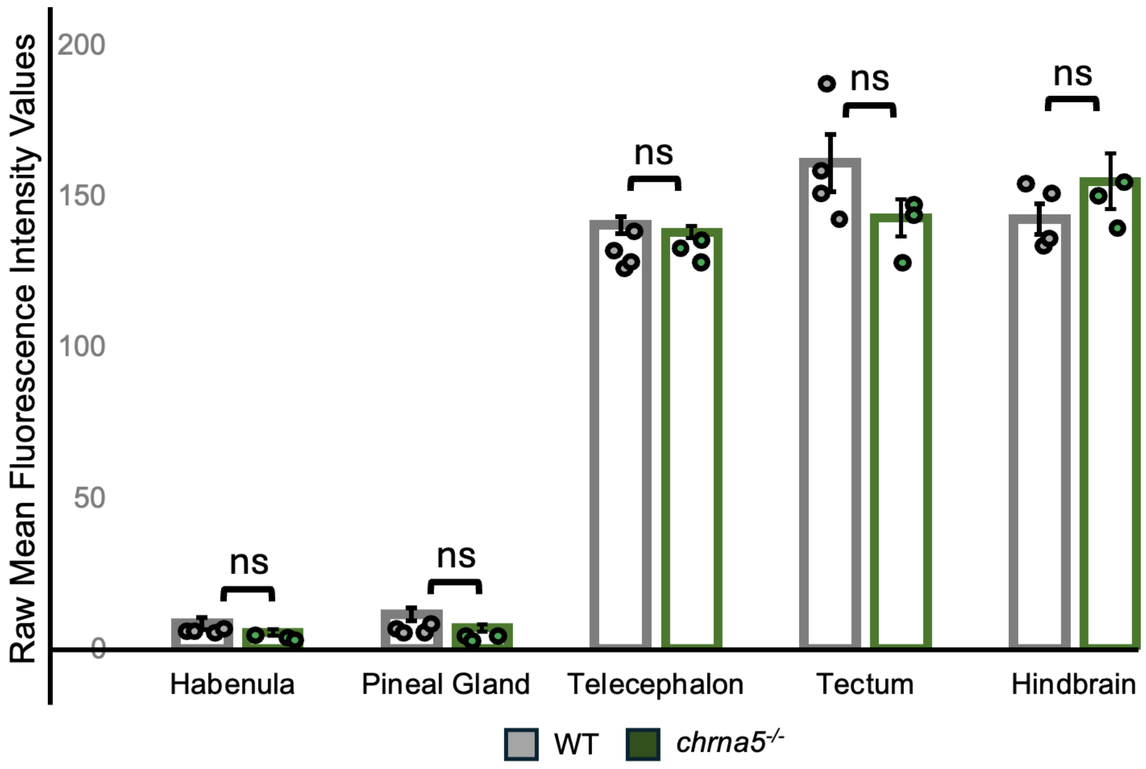

**S4. HCR quantitative data**

Values quantified and represented as bar plots from HCR images of both WT and chrna5$^{-/-}$ mutants, dorsally mounted to assess chrna5 gene expression across different brain regions in zebrafish.

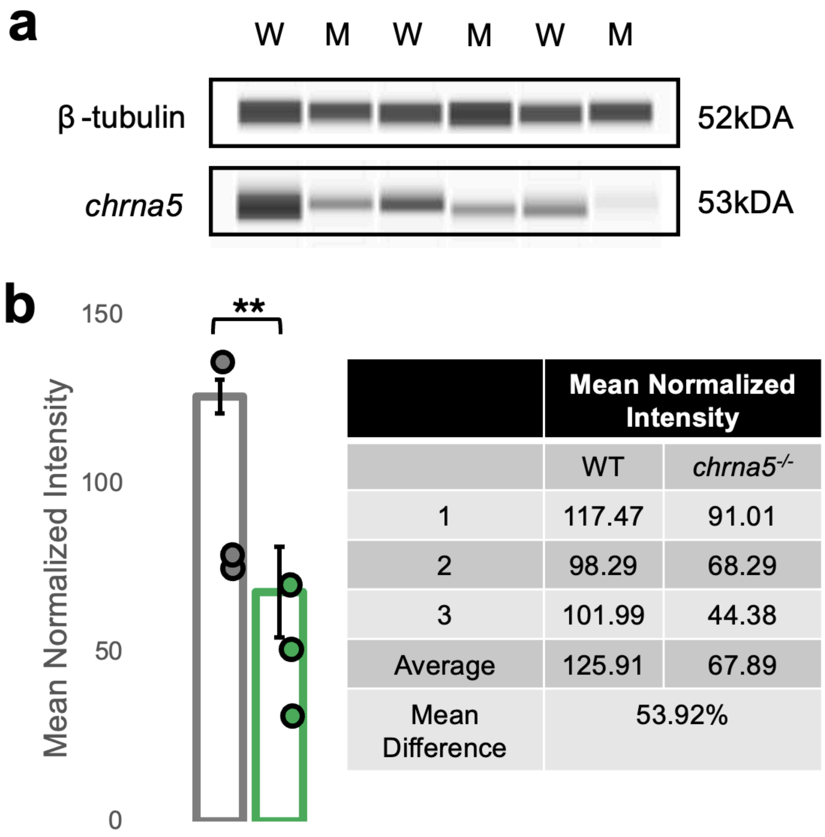

| | Mean Normalized Intensity | |
|---|---|---|
| | WT | *chrna5*$^{-/-}$ |
| 1 | 117.47 | 91.01 |
| 2 | 98.29 | 68.29 |
| 3 | 101.99 | 44.38 |
| Average | 125.91 | 67.89 |
| Mean Difference | 53.92% | |

**S5. Western blot analysis for WT and chrna5$^{-/-}$**

A) Western Blot displaying protein expression for chrna5$^{-/-}$ and β-tubulin for three biological replicates B) Averaged mean normalized intensity (against β-tubulin) for the three biological replicates. WT (W) chrna5 $^{-/-}$ M (M)

**Table S2.** Mean measured alcohol and predicted nicotine chamber concentrations during SAZA.

| Alcohol measured chamber concentrations | | | | | | |
|---|---|---|---|---|---|---|
| Time (min) | 70% mean | 70% SD | 40% mean | 40% SD | 20% mean | 20% SD |
| 0 | 0.00 | 0.00 | 0.00 | 0.00 | 0.00 | 0.00 |
| 1 | 0.70 | 0.00 | 0.47 | 0.06 | 0.30 | 0.10 |
| 2 | 0.80 | 0.28 | 0.37 | 0.06 | 0.33 | 0.06 |
| 3 | 1.15 | 0.35 | 0.70 | 0.17 | 0.40 | 0.00 |
| 4 | 1.35 | 0.21 | 0.60 | 0.17 | 0.50 | 0.10 |
| 5 | 1.50 | 0.28 | 0.73 | 0.23 | 0.47 | 0.06 |
| 6 | 1.50 | 0.14 | 0.70 | 0.12 | 0.50 | 0.00 |
| 7 | 1.75 | 0.21 | 0.73 | 0.06 | 0.53 | 0.06 |
| 8 | 1.85 | 0.21 | 0.90 | 0.36 | 0.57 | 0.15 |
| 9 | 1.95 | 0.07 | 0.90 | 0.26 | 0.60 | 0.10 |
| 10 | 1.85 | 0.07 | 0.93 | 0.15 | 0.63 | 0.06 |
| 11 | 1.85 | 0.07 | 0.83 | 0.21 | 0.67 | 0.06 |
| 12 | 1.95 | 0.21 | 0.77 | 0.12 | 0.70 | 0.00 |
| Mean ($\bar{x}$) | 1.52 | | 0.72 | | 0.52 | |
| **Alcohol flow rate adjustment factor** | | | | | | |
| Alcohol % | 0% | 20% | 40% | 70% | | |
| Flow rate per second (uL) | 27.74 | 16.94 | 11.32 | 10.61 | | |
| Flow normalisation factor (to 0%) | 1 | 1.63 | 2.45 | 2.61 | | |

$$\frac{Source\ concentration}{Flow\ normalisation\ factor \times alcohol\ mean\ measured\ chamber\ concentration} = Flow\ normalised\ dilution$$

| Predicted Nicotine chamber concentrations | | | | |
|---|---|---|---|---|
| Time (mins) | Mean flow normalised dilution factor | 500uM | 100uM | 10uM |
| 0 | NA | 0.00 | 0.00 | 0.00 |
| 1 | 38.07 | 13.13 | 2.63 | 0.26 |
| 2 | 38.29 | 13.06 | 2.61 | 0.26 |
| 3 | 25.77 | 19.40 | 3.88 | 0.39 |
| 4 | 23.87 | 20.94 | 4.19 | 0.42 |
| 5 | 22.15 | 22.58 | 4.52 | 0.45 |
| 6 | 21.91 | 22.82 | 4.56 | 0.46 |
| 7 | 20.20 | 24.75 | 4.95 | 0.50 |
| 8 | 18.10 | 27.63 | 5.53 | 0.55 |
| 9 | 17.45 | 28.66 | 5.73 | 0.57 |
| 10 | 17.12 | 29.20 | 5.84 | 0.58 |
| 11 | 17.50 | 28.57 | 5.71 | 0.57 |
| 12 | 17.53 | 28.53 | 5.71 | 0.57 |
| Mean ($\bar{x}$) | 23.16 | 23.27 | 4.65 | 0.47 |

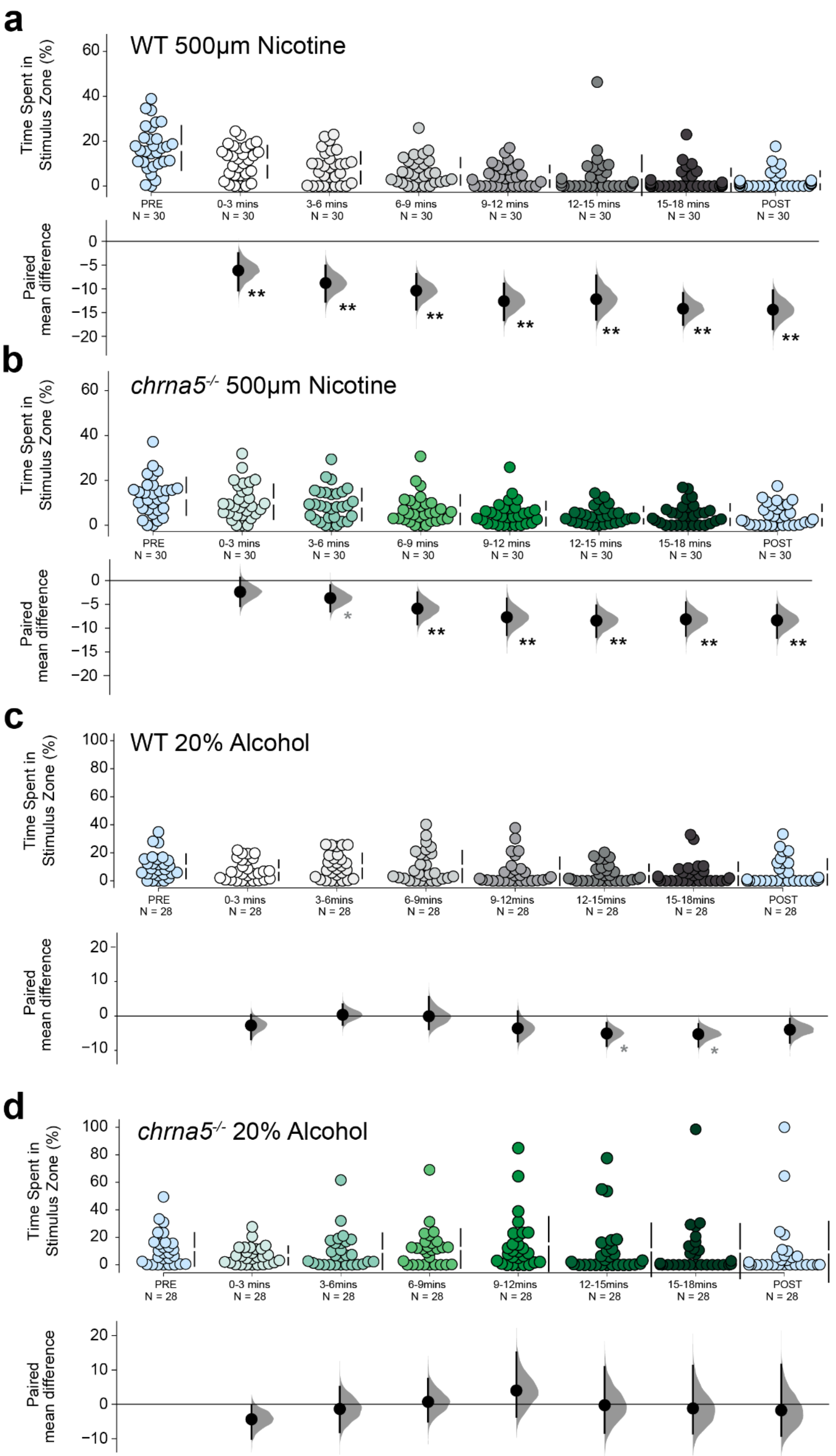

a
WT 500μm Nicotine
Time Spent in Stimulus Zone (%)
PRE N = 30
0-3 mins N = 30
3-6 mins N = 30
6-9 mins N = 30
9-12 mins N = 30
12-15 mins N = 30
15-18 mins N = 30
POST N = 30
Paired mean difference
b
chrna5-/- 500μm Nicotine
c
WT 20% Alcohol
PRE N = 28
0-3 mins N = 28
3-6mins N = 28
6-9mins N = 28
9-12mins N = 28
12-15mins N = 28
15-18mins N = 28
POST N = 28
d
chrna5-/- 20% Alcohol

**S6. Three-minute intervals for WT and chrna5 mutants**

A breakdown of the 18-minute stimulus period in three-minute intervals A) WT nicotine B) chrna5$^{-/-}$ mutants nicotine C) WT alcohol D) chrna5$^{-/-}$ mutants alcohol.

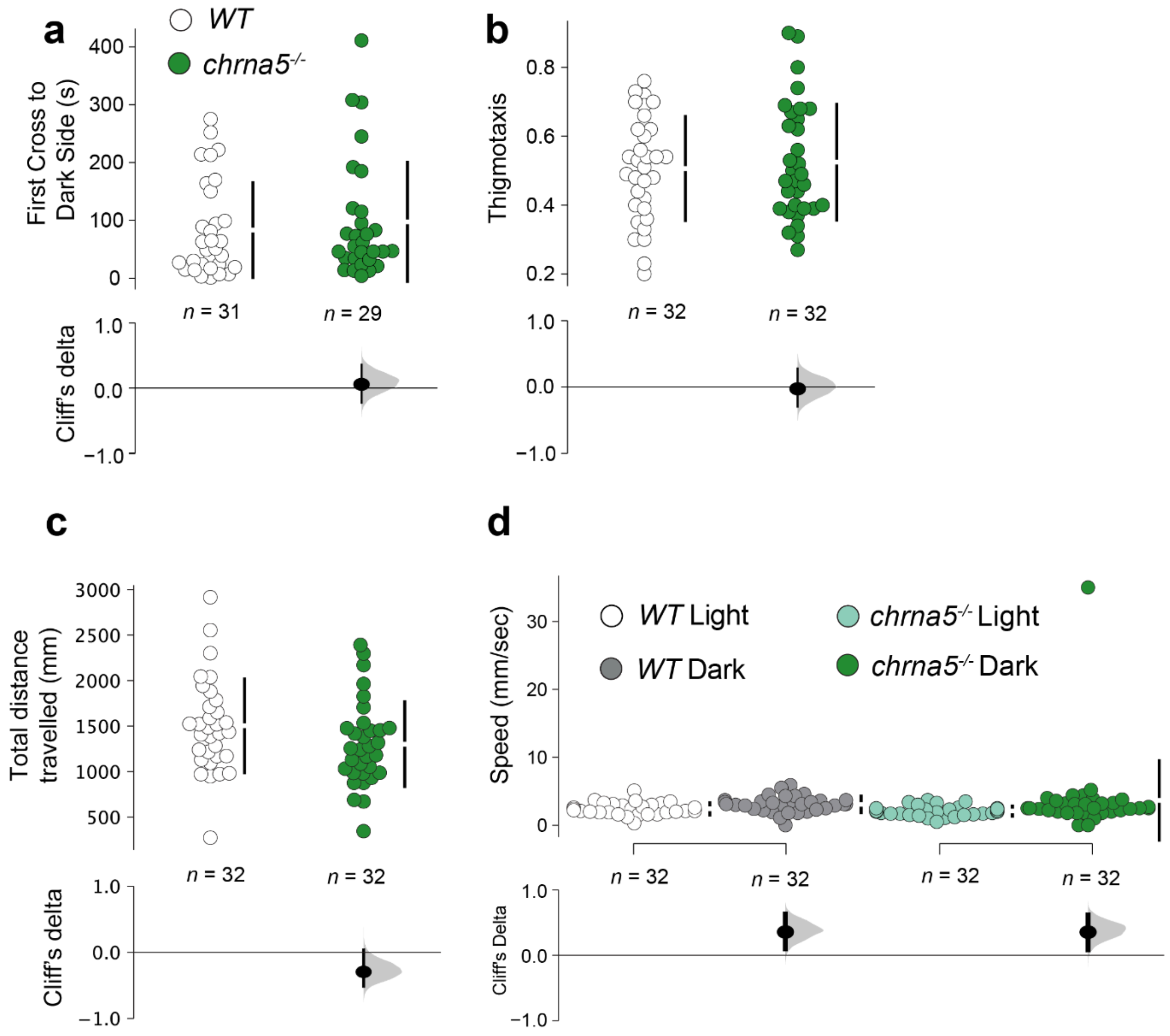

**S7. Anxiety graphs**

Displays additional graphs for WT and chrna5$^{-/-}$ mutants for A) total distance B) total speed for light and dark zones C) Total distance travelled D) Speed for WT light, dark and chrna5$^{-/-}$ mutants light, dark

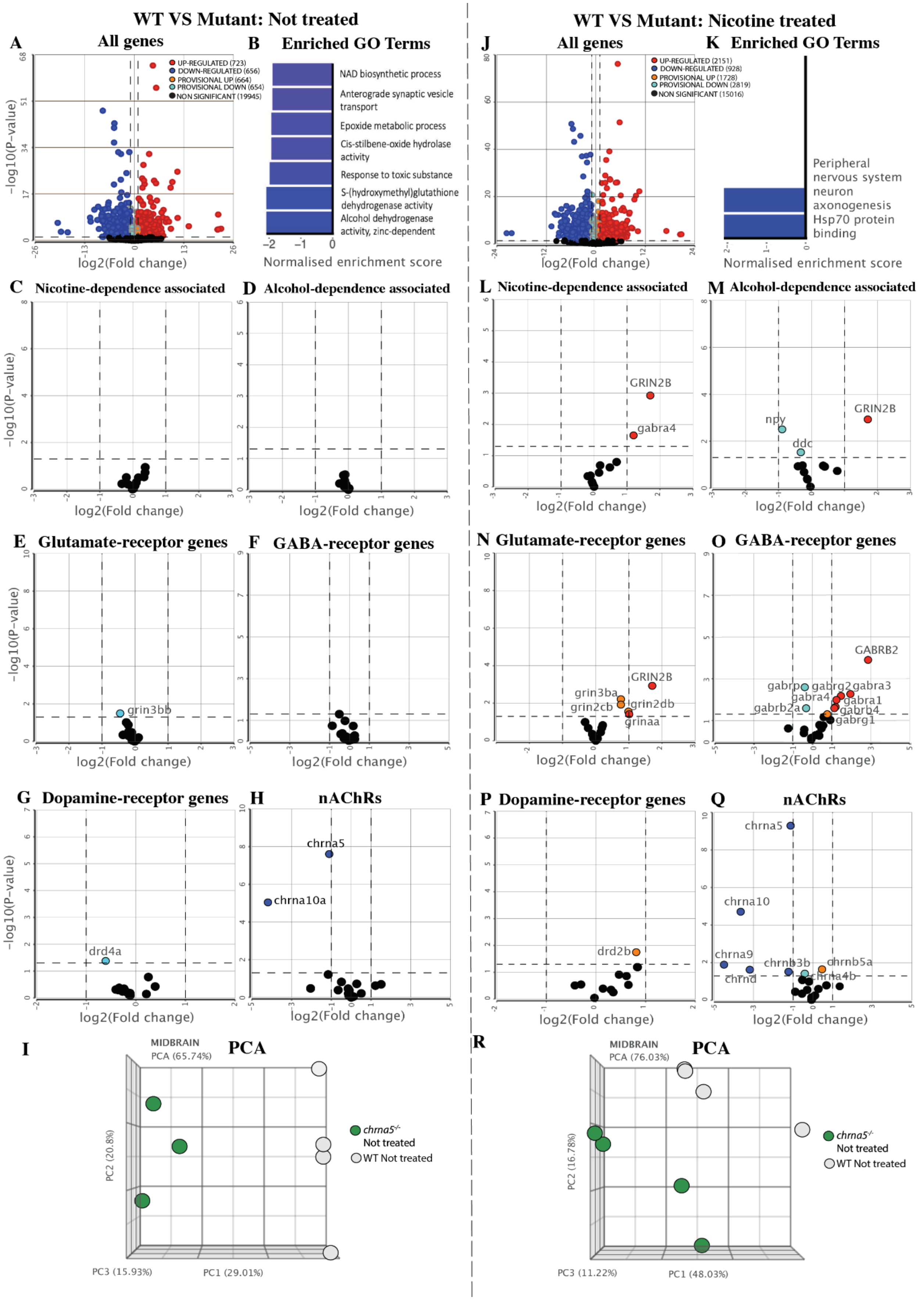
WT VS Mutant: Not treated
A All genes
B Enriched GO Terms
UP-REGULATED (723)
DOWN-REGULATED (656)
PROVISIONAL UP (664)
PROVISIONAL DOWN (654)
NON SIGNIFICANT (19945)
NAD biosynthetic process
Anterograde synaptic vesicle transport
Epoxide metabolic process
Cis-stilbene-oxide hydrolase activity
Response to toxic substance
S-(hydroxymethyl)glutathione dehydrogenase activity
Alcohol dehydrogenase activity, zinc-dependent
Normalised enrichment score
−log10(P-value)
log2(Fold change)
C Nicotine-dependence associated
D Alcohol-dependence associated
E Glutamate-receptor genes
grin3bb
F GABA-receptor genes
G Dopamine-receptor genes
drd4a
H nAChRs
chrna5
chrna10a
I PCA
MIDBRAIN
PCA (65.74%)
PC2 (20.8%)
PC3 (15.93%)
PC1 (29.01%)
chrna5-/- Not treated
WT Not treated
WT VS Mutant: Nicotine treated
J All genes
K Enriched GO Terms
UP-REGULATED (2151)
DOWN-REGULATED (928)
PROVISIONAL UP (1728)
PROVISIONAL DOWN (2819)
NON SIGNIFICANT (15016)
Peripheral nervous system neuron axonogenesis
Hsp70 protein binding
L Nicotine-dependence associated
GRIN2B
gabra4
M Alcohol-dependence associated
GRIN2B
npy
ddc
N Glutamate-receptor genes
GRIN2B
grin3ba
grin2cb
grin2db
grinaa
O GABA-receptor genes
GABRB2
gabrp
gabrg2
gabra3
gabra4
gabra1
gabrb2a
gabrb4
gabrg1
P Dopamine-receptor genes
drd2b
Q nAChRs
chrna5
chrna10
chrna9
chrnd
chrnb3b
chrnb5a
chrna4b
R PCA
MIDBRAIN
PCA (76.03%)
PC2 (16.78%)
PC3 (11.22%)
PC1 (48.03%)
chrna5-/- Not treated
WT Not treated

**S8. Volcano plots for nicotine treated vs non-treated**

A) WT and chrna5$^{-/-}$ mutants not nicotine treated for A) All genes in the genome B) Enriched GO plots C) Nicotine-dependent genes D) Alcohol-dependent genes E) Glutamate-receptor genes F) GABA-receptor genes G) Dopamine-receptor genes H) nAChRs genes I) PCA. WT and chrna5$^{-/-}$ mutants nicotine treated for J) All genes in the genome K) Enriched GO plots L) Nicotine-dependent genes M) Alcohol-dependent genes N) Glutamate-receptor genes O) GABA-receptor genes P) Dopamine-receptor genes Q) nAChRs genes R) PCA.

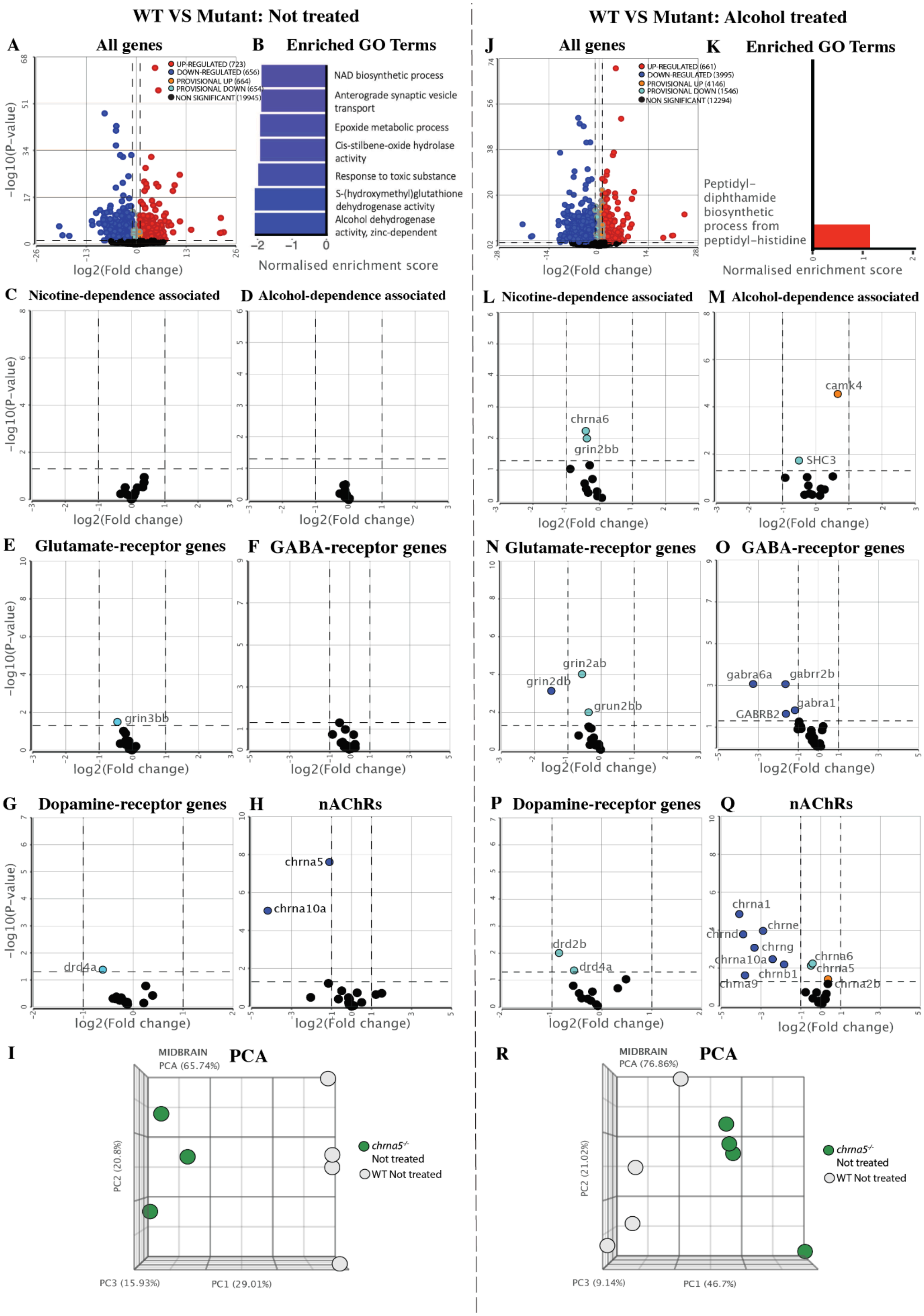
WT VS Mutant: Not treated
WT VS Mutant: Alcohol treated
A All genes
B Enriched GO Terms
UP-REGULATED (723)
DOWN-REGULATED (656)
PROVISIONAL UP (664)
PROVISIONAL DOWN (654)
NON SIGNIFICANT (19945)
NAD biosynthetic process
Anterograde synaptic vesicle transport
Epoxide metabolic process
Cis-stilbene-oxide hydrolase activity
Response to toxic substance
S-(hydroxymethyl)glutathione dehydrogenase activity
Alcohol dehydrogenase activity, zinc-dependent
Normalised enrichment score
J All genes
K Enriched GO Terms
UP-REGULATED (661)
DOWN-REGULATED (3995)
PROVISIONAL UP (4146)
PROVISIONAL DOWN (1546)
NON SIGNIFICANT (12294)
Peptidyl-diphthamide biosynthetic process from peptidyl-histidine
C Nicotine-dependence associated
D Alcohol-dependence associated
L Nicotine-dependence associated
M Alcohol-dependence associated
chrna6
grin2bb
camk4
SHC3
E Glutamate-receptor genes
F GABA-receptor genes
N Glutamate-receptor genes
O GABA-receptor genes
grin3bb
grin2ab
grin2db
grun2bb
gabra6a
gabrr2b
gabra1
GABRB2
G Dopamine-receptor genes
H nAChRs
P Dopamine-receptor genes
Q nAChRs
drd4a
chrna5
chrna10a
drd2b
chrna1
chrne
chrnd
chrng
chrnb1
chrna9
chrna2b
−log10(P−value)
log2(Fold change)
I PCA
R PCA
MIDBRAIN
PCA (65.74%)
PCA (76.86%)
PC2 (20.8%)
PC3 (15.93%)
PC1 (29.01%)
PC2 (21.02%)
PC3 (9.14%)
PC1 (46.7%)
chrna5-/- Not treated
WT Not treated

**S9. Volcano plots for alcohol treated vs non-treated**

A) WT and chrna5$^{-/-}$ mutants not alcohol treated for A) All genes in the genome B) Enriched GO plots C) Nicotine-dependent genes D) Alcohol-dependent genes E) Glutamate-receptor genes F) GABA-receptor genes G) Dopamine-receptor genes H) nAChRs genes I) PCA. WT and chrna5$^{-/-}$ mutants alcohol treated for J) All genes in the genome K) Enriched GO plots L) Nicotine-dependent genes M) Alcohol-dependent genes N) Glutamate-receptor genes O) GABA-receptor genes P) Dopamine-receptor genes Q) nAChRs genes R) PCA.

**Table S3.** Statistical analysis output table for qPCR

| Metric/Figure | Relative Expression (vs WT) | Cliff's Delta | 95% CI Upper | 95% CI Lower | P- Value |
|---|---|---|---|---|---|
| qPCR Relative Expression Figure 1C | *chrna5* | -0.438 | 0.156 | -0.875 | 0.1224 |
| qPCR Relative Expression Figure S1 | *chrna3* | -0.143 | 0.551 | -0.755 | 0.6092 |
| qPCR Relative Expression Figure S1 | *chrnb4* | -0.347 | 0.306 | -0.878 | 0.261 |

**Table S4**. Statistical analysis output table from SAZA Nicotine

| Metric/Figure | Nicotine concentration dispensed (vs 0uM) | Cliff's Delta | 95% CI Upper | 95% CI Lower | P - Value | Holm Bonferroni Adjusted P- Value |
|---|---|---|---|---|---|---|
| Preference Index WT Figure 2B | 10uM | -0.523 | -0.228 | -0.749 | 0.0004 | 0.0004 |
| | 500uM | -0.762 | -0.533 | -0.900 | <0.001 | <0.001 |
| Preference Index *chrna5*$^{-/-}$ Figure 2C | 10uM | 0.0724 | 0.363 | -0.239 | 0.6248 | 0.6248 |
| | 500uM | -0.166 | 0.143 | -0.448 | 0.2688 | 0.5376 |
| **Metric/Figure** | **NIcotine concentration dispensed (vs WT)** | **Cliff's Delta** | **95% CI Upper** | **95% CI Lower** | **P- Value** | **Holm Bonferroni Adjusted P- Value** |
| Preference Index WT Vs *chrna5*$^{-/-}$ | 0uM | -0.223 | 0.0793 | -0.497 | 0.1496 | NA |

| | | | | | | |
|---|---|---|---|---|---|---|
| comparisons Figure 2D | 10uM | 0.376 | 0.629 | 0.0644 | 0.013 | |
| | 500uM | 0.493 | 0.704 | 0.202 | 0.0012 | |
| **Metric/Figure** | **3-minute Time intervals (vs PRE)** | **Mean Difference** | **95% CI Upper** | **95% CI Lower** | **P- Value** | **Holm Bonferroni Adjusted P- Value** |
| WT Time Spent in Stimulus Zone 500uM Nicotine Figure S6 | 0-3 minutes | -6.154 | -2.532 | -10.341 | 0.0034 | 0.006 |
| | 3-6 minutes | -8.786 | -5.127 | -12.676 | 0.0002 | 0.0014 |
| | 6-9 minutes | -10.387 | -6.931 | -14.330 | <0.001 | 0.006 |
| | 9-12 minutes | -12.604 | -8.927 | -16.553 | <0.001 | 0.006 |
| | 12-15 minutes | -12.184 | -7.224 | -16.473 | <0.001 | 0.006 |
| | 15-18 minutes | -14.192 | -10.897 | -17.558 | <0.001 | 0.006 |
| | POST | -14.398 | -10.378 | -18.460 | <0.001 | 0.006 |
| *chrna5*$^{-/-}$ Time Spent in Stimulus Zone 500uM Nicotine Figure S6 | 0-3 minutes | -2.380 | 0.651 | -5.401 | 0.1346 | 0.1346 |
| | 3-6 minutes | -3.671 | -0.922 | -6.522 | 0.021 | 0.042 |
| | 6-9 minutes | -5.891 | -2.457 | -9.188 | 0.0026 | 0.0078 |
| | 9-12 minutes | -7.701 | -3.738 | -11.464 | 0.0006 | 0.003 |
| | 12-15 minutes | -8.425 | -5.210 | -11.876 | <0.001 | 0.004 |
| | 15-18 minutes | -8.142 | -4.553 | -11.621 | 0.0002 | 0.0014 |
| | POST | -8.362 | -5.0833 | -12.018 | 0.0004 | 0.0024 |
| **Metric/Figure** | **Parameter** | **Cliff's Delta** | **95% CI Upper** | **95% CI Lower** | **P- Value** | **Holm Bonferroni Adjusted P- Value** |
| Forest plot Preference Index | Preference Index | -0.762 | -0.533 | -0.9 | 0 | |
| | Entries into Stimulus Zone | -0.624 | -0.338 | -0.816 | 0 | |

| | | | | | | |
|---|---|---|---|---|---|---|
| 0uM vs 500uM WT Figure 2E | Time in Stimulus Zone | -0.738 | -0.507 | -0.882 | 0 | NA |
| | Mean time per Entry | -0.0222 | 0.271 | -0.324 | 0.881 | |
| | Mean Velocity | -0.5 | -0.202 | -0.729 | 0.0006 | |
| | Time in control Zone | -0.387 | -0.0689 | -0.624 | 0.0096 | |
| Forest plot Preference Index 0uM vs 500uM *chrna5*$^{-/-}$ Figure 2F | Preference Index | -0.166 | 0.143 | -0.448 | 0.2688 | NA |
| | Entries into Stimulus Zone | -0.0667 | 0.2448 | -0.368 | 0.6616 | |
| | Time in Stimulus Zone | -0.2667 | 0.0598 | -0.547 | 0.0774 | |
| | Mean time per Entry | -0.347 | -0.0322 | -0.602 | 0.021 | |
| | Mean Velocity | 0.320 | 0.579 | 0.00920 | 0.0374 | |
| | Time in control Zone | 0.195 | 0.483 | -0.108 | 0.2084 | |

**Table S5.** Statistical analysis output table from SAZA Alcohol

| **Metric/Figure** | **Alcohol concentration dispensed (vs 0%)** | **Cliff's Delta** | **95% CI Upper** | **95% CI Lower** | **P- Value** | **Holm Bonferroni Adjusted P- Value** |
|---|---|---|---|---|---|---|
| Preference Index WT Figure 2G | 5% | 0.0613 | 0.349 | -0.237 | 0.6822 | 0.6822 |
| | 10% | -0.226 | 0.113 | -0.514 | 0.1456 | 0.2912 |
| | 20% | -0.611 | -0.327 | -0.810 | <0.001 | <0.001 |
| Preference Index *chrna5*$^{-/-}$ | 5% | 0.447 | 0.691 | 0.121 | 0.0034 | 0.0068 |
| | 10% | 0.444 | 0.665 | 0.131 | 0.0018 | 0.0054 |

| Figure 2H | 20% | 0.109 | 0.399 | -0.181 | 0.4654 | 0.4654 |
|---|---|---|---|---|---|---|
| **Metric/Figure** | **Alcohol concentration dispensed (vs WT)** | **Cliff's Delta** | **95% CI Upper** | **95% CI Lower** | **P- Value** | **Holm Bonferroni Adjusted P- Value** |
| Preference Index WT Vs *chrna5*$^{-/-}$ Comparisons Figure 2I | 0% | -0.323 | -0.0258 | -0.577 | 0.0278 | NA |
| | 5% | 0.329 | 0.599 | 0.0104 | 0.0276 | |
| | 10% | 0.422 | 0.653 | 0.113 | 0.0056 | |
| | 20% | 0.430 | 0.667 | 0.135 | 0.0046 | |
| **Metric/Figure** | **3-minute Time intervals (vs PRE)** | **Mean Difference** | **95% CI Upper** | **95% CI Lower** | **P- Value** | **Holm Bonferroni Adjusted P- Value** |
| WT Time Spent in Stimulus Zone (%) 20% Alcohol Figure S6 | 0-3 minutes | -2.724 | 0.347 | -6.751 | 0.1314 | 0.468 |
| | 3-6 minutes | 0.362 | 3.363 | -2.580 | 0.8198 | 1 |
| | 6-9 minutes | -0.0750 | 5.608 | -3.853 | 0.9748 | 1 |
| | 9-12 minutes | -3.590 | 1.405 | -7.418 | 0.117 | 0.468 |
| | 12-15 minutes | -5.0592 | -1.977 | -8.744 | 0.0062 | 0.0372 |
| | 15-18 minutes | -5.259 | -2.337 | -8.895 | 0.0034 | 0.0238 |
| | POST | -3.973 | -0.766 | -7.803 | 0.0388 | 0.194 |
| *chrna5*$^{-/-}$ Time Spent in Stimulus Zone (%) 20% Alcohol Figure S6 | 0-3 minutes | -4.333 | -0.374 | -10.005 | 0.0966 | 0.6762 |
| | 3-6 minutes | -1.336 | 5.119 | -8.104 | 0.7124 | 1 |
| | 6-9 minutes | 0.728 | 7.444 | -5.0489 | 0.824 | 1 |
| | 9-12 minutes | 4.003 | 15.0888 | -3.614 | 0.4028 | 1 |
| | 12-15 minutes | -0.243 | 10.871 | -8.332 | 0.9638 | 1 |
| | 15-18 minutes | -1.178 | 11.232 | -8.476 | 0.8426 | 1 |
| | POST | -1.732 | 11.540 | -9.167 | 0.756 | 1 |

| Metric/Figure | Parameter | Cliff's Delta | 95% CI Upper | 95% CI Lower | P- Value | Holm Bonferroni Adjusted P- Value |
|---|---|---|---|---|---|---|
| Forest plot Preference Index 0% Vs 20% WT Figure 2J | Preference Index | -0.611 | -0.327 | -0.810 | 0 | NA |
| | Entries into Stimulus Zone | -0.164 | 0.167 | -0.473 | 0.2898 | |
| | Time in Stimulus Zone | -0.876 | -0.652 | -0.971 | 0 | |
| | Mean time per Entry | -0.981 | -0.914 | -1 | 0 | |
| | Mean Velocity | 0.860 | 0.957 | 0.583 | 0 | |
| | Time in control Zone | -0.771 | -0.507 | -0.919 | 0 | |
| Forest plot 0% Vs 20% *chrna5*$^{-/-}$ Figure 2K | Preference Index | 0.109 | 0.399 | -0.181 | 0.4654 | NA |
| | Entries into Stimulus Zone | -0.158 | 0.153 | -0.445 | 0.2884 | |
| | Time in Stimulus Zone | -0.240 | 0.0691 | -0.521 | 0.111 | |
| | Mean time per Entry | -0.237 | 0.0899 | -0.530 | 0.1132 | |
| | Mean Velocity | 0.754 | 0.893 | 0.484 | 0 | |
| | Time in control Zone | -0.675 | -0.401 | -0.850 | 0 | |

**Table S6.** Statistical analysis output table for pretreatment Nicotine

| Metric/Figure | Parameter | Cliff's Delta | 95% CI Upper | 95% CI Lower | P- Value |
|---|---|---|---|---|---|
| Pretreatment Nicotine/Nicotine SA Forest plot Preference Index 0uM Vs 500uM WT Figure 4A | Preference index | -0.792 | -0.572 | -0.917 | <0.001 |
| | Entries into Stimulus Zone | -0.739 | -0.521 | -0.878 | <0.001 |
| | Time in Stimulus Zone | -0.867 | -0.699 | -0.953 | <0.001 |
| | Mean time per Entry | -0.434 | -0.123 | -0.673 | 0.0028 |
| | Mean Velocity | 0.174 | 0.458 | -0.144 | 0.2366 |
| | Time in control zone | -0.0731 | 0.230 | -0.368 | 0.6254 |
| Pretreatment Nicotine/Nicotine SA Forest plot Preference Index 0uM Vs 500uM *chrna5*$^{-/-}$ Figure 4B | Preference index | -0.414 | -0.0989 | -0.667 | 0.0058 |
| | Entries into Stimulus Zone | -0.292 | 0.00345 | -0.567 | 0.049 |
| | Time in Stimulus Zone | -0.541 | -0.230 | -0.757 | <0.001 |
| | Mean time per Entry | -0.489 | -0.189 | -0.717 | 0.0008 |
| | Mean Velocity | 0.282 | 0.553 | -0.0230 | 0.0668 |
| | Time in control zone | 0.322 | 0.584 | 0.00575 | 0.0346 |
| Preference Index WT Vs *chrna5*$^{-/-}$ Comparisons Figure 4C | Acute | 0.493 | 0.704 | 0.202 | 0.0012 |
| | Pretreatment nicotine/nicotine SA | 0.338 | 0.578 | 0.0258 | 0.0244 |
| Forest plot Preference Index | Preference index | -0.383 | -0.0713 | -0.637 | 0.011 |
| | Entries into Stimulus Zone | 0.0598 | 0.384 | -0.263 | 0.6864 |
| | Time in Stimulus Zone | -0.795 | -0.591 | -0.908 | <0.001 |

| | | | | | |
|---|---|---|---|---|---|
| 0uM Vs 500uM Pretreatment nicotine/alcohol SA WT Figure 4D | Mean time per Entry | -0.839 | -0.641 | -0.940 | <0.001 |
| | Mean Velocity | 0.949 | 0.986 | 0.828 | <0.001 |
| | Time in control zone | -0.959 | -0.855 | -0.993 | <0.001 |
| Forest plot Preference Index 0uM Vs 500uM Pretreatment nicotine/alcohol SA *chrna5*$^{-/-}$ Figure 4E | Preference index | 0.335 | 0.587 | 0.0516 | 0.0252 |
| | Entries into Stimulus Zone | -0.0333 | 0.271 | -0.326 | 0.8194 |
| | Time in Stimulus Zone | -0.0688 | 0.237 | -0.355 | 0.6414 |
| | Mean time per Entry | -0.135 | 0.191 | -0.426 | 0.3692 |
| | Mean Velocity | 0.775 | 0.9 | 0.5412 | <0.001 |
| | Time in control zone | -0.785 | -0.4903 | -0.929 | <0.001 |
| Preference Index WT Vs *chrna5*$^{-/-}$ Comparisons Figure 4F | Acute | 0.402 | 0.640 | 0.110 | 0.008 |
| | Pretreatment nicotine/alcohol SA | 0.408 | 0.637 | 0.117 | 0.0058 |

**Table S7.** Statistical analysis output table from pretreatment Alcohol

| Metric/Figure | Parameter | Cliff's Delta | 95% CI Upper | 95% CI Lower | P- Value |
|---|---|---|---|---|---|
| Forest plot Preference Index 0% Vs 20% WT Figure 5A | Preference index | -0.617 | -0.347 | -0.799 | <0.001 |
| | Entries into Stimulus Zone | -0.220 | 0.0760 | -0.496 | 0.1332 |
| | Time in Stimulus Zone | -0.931 | -0.717 | -0.994 | <0.001 |
| | Mean time per Entry | -0.931 | -0.738 | -0.990 | <0.001 |

| | | | | | |
|---|---|---|---|---|---|
| | Mean Velocity | 0.960 | 0.990 | 0.858 | <0.001 |
| | Time in control zone | -0.929 | -0.775 | -0.990 | <0.001 |
| Forest plot Preference Index 0% Vs 20% *chrna5*$^{-/-}$ Figure 5B | Preference index | -0.0278 | 0.294 | -0.337 | 0.862 |
| | Entries into Stimulus Zone | -0.206 | 0.101 | -0.478 | 0.167 |
| | Time in Stimulus Zone | -0.0478 | 0.257 | -0.339 | 0.7562 |
| | Mean time per Entry | 0.168 | 0.466 | -0.150 | 0.2776 |
| | Mean Velocity | 0.5 | 0.739 | 0.181 | 0.0004 |
| | Time in control zone | -0.495 | -0.181 | -0.724 | 0.0014 |
| Preference Index WT Vs *chrna5*$^{-/-}$ Comparisons Figure 5C | Acute | 0.401 | 0.640 | 0.110 | 0.008 |
| | Pretreatment alcohol/alcohol SA | 0.225 | 0.504 | -0.0927 | 0.1302 |
| Forest plot Preference Index 0% Vs 20% Pretreatment alcohol/nicotine SA WT Figure 5D | Preference index | -0.837 | -0.602 | -0.95 | <0.001 |
| | Entries into Stimulus Zone | -0.746 | -0.493 | -0.896 | <0.001 |
| | Time in Stimulus Zone | -0.924 | -0.767 | -0.979 | <0.001 |
| | Mean time per Entry | -0.629 | -0.331 | -0.824 | <0.001 |
| | Mean Velocity | 0.502 | 0.740 | 0.219 | 0.0006 |
| | Time in control zone | -0.0690 | 0.243 | -0.374 | 0.649 |
| Forest plot | Preference index | -0.502 | -0.182 | -0.731 | 0.0014 |
| | Entries into Stimulus Zone | -0.660 | -0.369 | -0.844 | <0.001 |

| | | | | | |
|---|---|---|---|---|---|
| Preference Index 0% Vs 20% Pretreatment alcohol/nicotine SA *chrna5*$^{-/-}$ Figure 5E | Time in Stimulus Zone | -0.599 | -0.319 | -0.803 | <0.001 |
| | Mean time per Entry | -0.128 | 0.191 | -0.427 | 0.4038 |
| | Mean Velocity | -0.114 | 0.218 | -0.394 | 0.4654 |
| | Time in control zone | 0.0916 | 0.386 | -0.227 | 0.5422 |
| Preference Index WT Vs *chrna5*$^{-/-}$ Comparisons Figure 5F | Acute | 0.493 | 0.704 | 0.202 | 0.0012 |
| | Pretreatment alcohol/nicotine SA | 0.260 | 0.527 | -0.0603 | 0.0974 |

**Table S8**. Statistical analysis output table for the comorbid disorders

| **Metric/Figure** | **Parameter** | **Cliff's Delta / Paired mean difference** | **95% CI Upper** | **95% CI Lower** | **P- Value** | **Holm Bonferroni Adjusted P- Value** |
|---|---|---|---|---|---|---|
| Anxiety WT Vs *chrna5*$^{-/-}$ | Time spent in dark zone Figure 6B | -0.0479 | 0.239 | -0.333 | 0.7404 | NA |
| | Number of Entries into Dark side Figure 6B | -0.139 | -0.139 | -0.407 | 0.3342 | |
| | First Cross Dark side Figure S7A | 0.0857 | 0.358 | -0.224 | 0.5616 | |
| | Thigmotaxis Figure S7B | 0.000977 | 0.278 | -0.295 | 0.9928 | |
| | Distance Figure S7C | -0.268 | 0.0391 | -0.514 | 0.0642 | |
| | Velocity | 0.384 | 0.627 | 0.0986 | 0.0066 | |

| | | | | | | |
|---|---|---|---|---|---|---|
| | (mm/sec) Light Figure S7D | | | | | |
| | Velocity (mm/sec) Dark Figure S7D | 0.382 | 0.617 | 0.0840 | 0.0082 | |
| Circadian Rhythm WT Figure 6F | Night | 0.514 | 0.690 | 0.284 | <0.001 | <0.001 |
| | After Night | -0.797 | -0.635 | -0.894 | <0.001 | <0.001 |
| Circadian Rhythm *chrna5*$^{-/-}$ MUT Figure 6G | Night | 0.840 | 0.922 | 0.699 | <0.001 | <0.001 |
| | After Night | -0.703 | -0.514 | -0.834 | <0.001 | <0.001 |
| Circadian Rhythm WT Figure 6K | Light -> Dark transition WT | 44 | 48.7 | 39.4 | < 0.001 | |
| Circadian Rhythm *chrna5*$^{-/-}$ MUT Figure 6K | Light -> Dark transition *chrna5*$^{-/-}$ MUT | 27.9 | 33.3 | 25.9 | < 0.001 | |
| Circadian Rhythm Intergenotype Figure 6K | Light -> Dark transition delta-delta | -14.2 | -8.78 | -19.9 | 0.0026 | NA |
| Circadian Rhythm WT Figure 6K | Dark -> Light transition WT | 13.7 | 16 | 11.6 | < 0.001 | |
| Circadian Rhythm | Dark -> Light transition | 9.44 | 10.7 | 8.1 | < 0.001 | |

| | | | | | | |
|---|---|---|---|---|---|---|
| *chrna5*$^{-/-}$ MUT Figure 6L | *chrna5*$^{-/-}$ MUT | | | | | |
| Circadian Rhythm Intergenotype Figure 6L | Dark -> Light transition delta-delta | -4.25 | -1.67 | -6.79 | 0.014 | |
| Appetite WT Vs *chrna5*$^{-/-}$ Egg Yolk Figure 6N | Mean Fluorescent Intensity | -0.203 | -0.109 | -0.296 | 0 | NA |
| Appetite WT Vs *chrna5*$^{-/-}$ Protein Figure 6O | Mean Fluorescent Intensity | 0.312 | 0.218 | 0.404 | 0 | NA |
| Appetite WT Vs *chrna5*$^{-/-}$ Paramecium Figure 6P | Mean Fluorescent Intensity | 0.180 | 0.274 | 0.088 | <0.001 | NA |

**Table S9.** Statistical analysis output table for vibration assay Figure 6I, linear mixed effects model with repeated measures between genotype per intensity.

| Metric/Figure | Intensity | Repeated measures estimated marginal mean difference | Standard Error | Z Ratio | P- Value |
|---|---|---|---|---|---|
| Circadian Rhythm vibration assay (Fig 6I) | 0 | -1.25 | 7.90 | -0.16 | 0.873841 |
| | 0.12 | 1.47 | 7.90 | 0.19 | 0.852319 |
| | 0.24 | 5.57 | 7.90 | 0.70 | 0.481011 |
| | 0.36 | 11.09 | 7.90 | 1.40 | 0.160614 |
| | 0.48 | 14.87 | 7.90 | 1.88 | 0.059824 |
| | 0.6 | 21.36 | 7.90 | 2.70 | 0.006864 |
| | 0.72 | 25.37 | 7.90 | 3.21 | 0.001325 |
| | 0.84 | 32.10 | 7.90 | 4.06 | 0.000049 |

| | | | | | |
|---|---|---|---|---|---|
| | 1 | 39.25 | 7.90 | 4.97 | 0.000001 |

**Table S10.** Genes included in the custom sets ‘Nicotine dependence ’ and ‘Alcohol dependence’. These genes have been associated with nicotine and alcohol dependence, described by Hu *et al.* (Hu et al. 2018)

| Gene set | Gene list |
|---|---|
| Nicotine dependence | grin1a<br>gabra4<br>grin2aa<br>grin2bb<br>GRIN2B<br>GABRA2A<br>chrnb2<br>chrnb5b<br>chrna6<br>chrna7<br>FO907089.1 |
| Alcohol dependence | camk4<br>ppp1r1b<br>ddc<br>gnas<br>grin3a<br>npy<br>bdnf<br>grin2b<br>shc3<br>th<br>slc6a3<br>slc18a2 |
